# History of previous midlife estradiol treatment permanently alters interactions of brain insulin-like growth factor-1 signaling and hippocampal estrogen synthesis to enhance cognitive aging in a rat model of menopause

**DOI:** 10.1101/2022.03.31.486639

**Authors:** Nina E. Baumgartner, Shannon M. McQuillen, Samantha F. Perry, Sangtawan Miller, Robert Gibbs, Jill M Daniel

## Abstract

Across species, including humans, elevated levels of brain estrogen receptor (ER) α are associated with enhanced cognitive aging even in the absence of circulating estrogens. In rodents, short-term estrogen treatment—such as that commonly used in the menopausal transition—results in long-term increases in levels of ERα in the hippocampus, leading to enhanced memory long after termination of estrogen treatment. However, mechanisms by which increased levels of brain ERα enhances cognitive aging remain unclear. Here we show that in the hippocampus, insulin-like growth factor-1 (IGF-1)—which can activate ER via ligand-independent mechanisms—requires concomitant synthesis of brain-derived neuroestrogens to phosphorylate ERα via MAPK signaling, ultimately resulting in enhanced memory. In a rat model of menopause involving long-term ovarian hormone deprivation, hippocampal neuroestrogen activity decreases, altering IGF-1 activity and resulting in impaired memory. However, this process is reversed by short-term estradiol treatment. Forty-days of estradiol exposure following ovariectomy results in maintenance of neuroestrogen levels that persist beyond the period of hormone treatment, allowing for continued interactions between IGF-1 and neuroestrogen signaling, elevated levels of hippocampal ERα, and ultimately enhanced memory. Collectively, results demonstrate that short-term estradiol use following loss of ovarian function has long-lasting effects on hippocampal function and memory by dynamically regulating cellular mechanisms that promote activity of ERα in the absence of circulating estrogens. Translational impacts of these findings suggest lasting cognitive benefits of short-term estrogen use near menopause and highlight the importance of hippocampal ERα—independent from the role of circulating estrogens—in regulating memory in aging females.

**Significance statement:** Declines in ovarian hormones following menopause coincide with increased risk of cognitive decline. Due to potential health risks, current recommendations are that menopausal estrogen therapy be limited to a few years. Long-term consequences for the brain and memory of this short-term midlife estrogen therapy are unclear. Here, in a rodent model of menopause, we determined mechanisms by which short-term midlife estrogen exposure can enhance hippocampal function and memory with cognitive benefits and molecular changes enduring long after termination of estrogen exposure. Our model indicates long-lasting benefits of maintaining hippocampal estrogen receptor function in the absence of ongoing estrogen exposure and suggests potential strategies for combating age-related cognitive decline.

## Introduction

Loss of ovarian hormones during menopause coincides with cognitive decline and increased risk of age-related dementias (Henderson et al. 1996; Sherwin 1994). Due to putative health risks associated with prolonged estrogen exposure, current health guidelines recommend using menopausal estrogen treatment for as short a time as possible. Work from our lab in a rodent model of menopause has demonstrated long-lasting benefits of short-term midlife estradiol treatment on hippocampal function and memory through sustained activation of estrogen receptor (ER) α that are likely permanent, persisting long after estradiol treatment is terminated (Rodgers et al 2010; Witty et al 2013; Black et al. 2016; Baumgartner et al. 2021). These findings correspond with evidence across multiple species, including humans, that elevated levels of brain estrogen receptor ERα are associated with enhanced cognitive aging even in the absence of circulating estrogens (For review, see Baumgartner and Daniel, 2020). The mechanisms by which increased levels of brain ERα enhance cognitive aging following previous midlife exposure to estradiol are unclear.

Short-term exposure to estradiol in midlife enhances memory and increases levels of hippocampal ERα long-term in ovariectomized rodents (Rodgers et al. 2010), effects dependent on insulin-like growth factor-1 (IGF-1) signaling (Witty et al. 2013) and resulting in sustained ER-dependent transcriptional activity (Pollard et al. 2018). IGF-1 is a peptide hormone that acts through IGF-1R, a tyrosine kinase receptor with much functional overlap with ERα, including activation of MAPK and PI3K-AKT signaling pathways by both receptors (Russo et al. 2005; Sohrabji 2015). ERα and IGF-1R co-localize and form estradiol-dependent protein complexes in the hippocampus (Cardona-Gomez et al. 2000; Mendez et al. 2003). Implications for these subcellular interactions for cognition remain to be determined. IGF-1 administration activates ERα via ligand-independent mechanisms *in vitro* (Kato et al. 1995) and in recently ovariectomized rats via phosphorylation at serine-118 (Grissom and Daniel 2016), a phospho-site crucial for protecting ERα from degradation (Valley et al. 2005). An *in vitro* study revealed that neuroestrogen synthesis is required for IGF-1-mediated activation of ERα, potentially through synergistic activation of the MAPK pathway (Pollard and Daniel 2019).

Contradicting a potential role for neuroestrogens in activation of ERα following loss of ovarian function are data indicating that hippocampal neuroestrogens are regulated by circulating estrogens (Nelson et al. 2016) and demonstrations of decreases in hippocampal aromatase expression and neuroestrogen levels following long-term ovariectomy (Chen et al. 2021; Ma et al 2020). Additionally, we showed that blocking neuroestrogen synthesis via aromatase inhibition had no impact on hippocampal ER-dependent transcription in long-term ovariectomized mice (Baumgartner et al 2019). Collectively, these data point to a diminished role for neuroestrogen synthesis in hippocampal function following long-term ovarian hormone deprivation.

In summary, data indicate that a history of midlife estradiol treatment impacts memory long after termination of estradiol treatment through lasting activation of hippocampal ERα by ligand-independent mechanisms via IGF1-signaling. *In vitro* evidence indicates that ligand-independent activation of ER by IGF-1 requires concomitant neuroestrogen synthesis. However, neuroestrogen levels in the hippocampus are diminished following long-term loss of ovarian hormones. The goal of the current work was to reconcile these contradictory findings and determine implications for female cognitive aging following loss of ovarian function of the interactive actions of IGF-1 and neuroestrogens and determine if history of estradiol use impacts that interaction. First, we determined the necessity of neuroestrogen synthesis in the ability of IGF-1 to activate ERα *in vivo* via its downstream signaling pathways and subsequent impact on memory. Next, we determined if interactions of neuroestrogens and IGF-1 in the hippocampus and subsequent impact on memory were altered in two models of menopause— one with and one without a history of past midlife estradiol use. Our findings provide a potential model for combatting postmenopausal cognitive decline in which short-term estradiol treatment following the loss of ovarian hormones sustains hippocampal function and memory well beyond the period of estradiol exposure by permanently altering the dynamic relationship between IGF-1R signaling and neuroestrogen synthesis.

## Materials and Methods

### Subjects

Middle-aged female Long-Evans hooded rats (Envigo), retired breeders (∼11 months of age), were used for all experiments. Animal care was in accordance with guidelines set by the National Institute of Health Guide for the Care and Use of Laboratory Animals and all procedures were approved by the Institutional Animal Care and Use Committee of Tulane University. Rats were housed individually in a temperature-controlled vivarium under a 12-h light, 12-h dark cycle and had unrestricted access to food and water unless otherwise noted.

### Ovariectomies and hormone treatments

All rats in experiments were anesthetized by intraperitoneal injections of ketamine (100 mg/kg ip; Bristol Laboratories, Syracuse, NY) and xylazine (7 mg/kg ip; Miles Laboratories, Shawnee, KS) and ovariectomized. Buprenorphine (0.375 mg/kg; Reckitt Benckiser Health Care) was administered by subcutaneous injection before the start of each surgery. Ovariectomy surgery involved bilateral flank incisions through the skin and muscle wall and the removal of ovaries.

For Experiments 3 and 4, rats were implanted with a subcutaneous 5-mm SILASTIC brand capsule (0.058 in. inner diameter and 0.077 in. outer diameter; Dow Corning, Midland, MI) on the dorsal aspect of the neck immediately following ovariectomy. Capsules contained either cholesterol vehicle (Experiment 3; Sigma-Aldrich, St. Louis, MO) or 25% 17β-estradiol (Experiment 4; Sigma-Aldrich) diluted in vehicle. We have previously shown that implants of these dimensions and estradiol concentrations maintain blood serum estradiol levels in middle-age retired breeders at approximately 37 pg/mL (Bohacek & Daniel 2007), which falls within physiological range. Forty days after ovariectomy and capsule implantation, capsules were removed. Vaginal smears for each rat were collected for at least four consecutive days before capsule replacement in order to confirm hormone treatment for the initial forty-day window. Smears of ovariectomized, cholesterol-treated rats were characterized by a predominance of leukocytes, while smears of ovariectomized, estradiol-treated rats were characterized by a predominance of cornified and nucleated epithelial cells.

### Stereotaxic surgeries

Rats were anesthetized with ketamine and xylazine as described above and administered buprenorphine as an analgesic. Rats were then placed into a stereotaxic frame. An incision was made in the scalp and fascia that overlie the skull and a hole was drilled in the skull.

In Experiment 1, a cannula connected to a Hamilton syringe via silastic tubing was lowered through the hole to the appropriate depth to reach the right lateral ventricle (relative to bregma: anteroposterior, −0.5 mm; mediolateral, −1.1 mm; dorsoventral, −2.5 mm). Cannulas delivered 5 uL of either vehicle containing 8% DMSO (Sigma-Aldrich) in aCSF (Tocris), 2 ug of human IGF-1 (GroPep) diluted in vehicle, or 2ug of IGF-1 combined with 0.4 ug of aromatase inhibitor Letrozole (Sigma-Aldrich) diluted in vehicle over the course of 10 minutes.

In Experiments 2-4, a cannula (brain infusion kits, Alzet) was lowered through the hole to the appropriate depth to reach the right lateral ventricle (relative to bregma: anteroposterior, −0.3 mm; mediolateral, −1.2 mm; dorsoventral, −4.5 mm) and adhered to the skull with an anchoring screw, Super Glue, and dental acrylic. The cannula was connected to an osmotic mini-pump (flow rate, 0.15 μl/h; Alzet) by vinyl tubing for drug delivery. Rats in Experiment 2 received mini-pumps that delivered either vehicle containing 6.7% DMSO in aCSF, human IGF-1 (0.33 ug/ul) diluted in vehicle, or human IGF-1 (0.33 ug/ul) and letrozole (0.066 ug/ul) diluted in vehicle. Rats from Experiments 3 and 4 received mini-pumps that delivered either vehicle (8% DMSO in aCSF), IGF-1 receptor antagonist JB1 in vehicle (300 μg/mL, Bachem), aromatase inhibitor letrozole in vehicle (0.066 ug/ul), or both JB1 + letrozole in vehicle.

### Radial-arm maze training

Approximately one week before the start of behavioral training, rats were food restricted and weighed daily to maintain their body weights at 85-90% of their free-feeding weight. Rats then began training on the 8-arm radial-maze task (Coulbourn Instruments, Whitehall, PA), as previously described (Daniel 2015). The maze consists of eight arms (66 cm long × 9.5 cm wide × 11.5 cm high) with a metal grated floor and clear acrylic walls. Arms extend out radially from a central hub that is 28 cm in diameter and the maze was placed on a table that is approximately 1 m above the ground. The maze was centered in a 3 × 5 m room with many visible extra maze cues. During training, a single food reward (Froot Loops; Kellogg Co., Battle Creek, MI) was placed in an opaque dish, 5.5 cm in diameter and 1.25 cm tall, at the end of each arm so it was not visible from the center of the maze. For each trial, the rat was placed in the center of the maze facing one of the eight arms. The starting orientation varied pseudo-randomly across trials. The rat was then allowed to enter arms and obtain food rewards until all eight arms had been visited or five minutes had elapsed. An arm entry was scored when all four paws crossed the midline of the arm. The arm entry sequence was scored in real time by an observer located in a fixed location in the room. Errors were scored if the rat re-entered an arm that had already been visited previously in the trial. Rats were trained with one trial per day, five days per week, for up to twenty-five days until they reached criterion by scoring fewer than 2 errors for three consecutive days. Once criterion was reached, rats underwent stereotaxic surgery and drug delivery as described above.

### Delay testing on radial-arm maze

One week after stereotaxic surgeries, rats were tested on delay trials. During testing, delays of various lengths were imposed which required the rats to remember over an extended period of time which arms had previously been visited. Rats were placed in the center of the maze facing one of the eight arms and allowed to enter four unique arms during the pre-delay trial. After four correct arm choices, rats were removed from the maze and placed in a holding cage for the duration of the delay. Following the delay, rats were returned to the center of the maze in the same orientation from the pre-delay trial. During this post-delay trial, rats were allowed to explore the maze until the four remaining, still baited arms were visited or until 5 minutes had elapsed. Re-entries into previously visited arms were recorded as errors. Arm-choice accuracy was measured by errors of eight, which represented the number of errors included in the first eight arm choices collectively across the pre-delay and post-delay trials. Rats received 2 days of habituation to a 1-min delay trial. Each subsequent delay was tested across two consecutive days. Delays for each experiment were chosen based on the performance of the rats during the training period and were increased in difficulty until at least two experimental groups performed within one standard deviation from chance (2.7 errors of eight). Means of errors of eight across both days of testing for each delay were analyzed.

### Euthanasia and tissue collection

Rats were killed under anesthesia induced by ketamine and xylazine. The hippocampus was dissected out and quick frozen on dry ice and stored at −80°C until further processing. A 1-cm sample of the right uterine horn was collected from each rat and weighed to verify ovariectomy status or hormone treatment at the time of euthanasia.

### Tissue processing and western blotting

In Experiment 1, right hippocampi were processed for subcellular protein fractionation and compartment-specific western blotting as previously described (Baumgartner et al. 2021). Briefly, hippocampal tissue was homogenized using the PowerGen-125 handheld homogenizer (Fisher), strained through Pierce Tissue Strainers, and separated into cytosolic, membrane, and nuclear compartments using consecutive centrifugation steps of varying speeds and specialized buffers obtained from a commercially available kit (Sub-Cellular Protein Fractionation Kit for Tissues, ThermoFisher). Bradford protein assays were performed for each compartment individually. Each sample diluted 1:1 with Laemmli Sample Buffer (BioRad) mixed with 350 mM DTT, boiled for 5 min, and stored at −80°C until western blotting.

The left hippocampi in Experiment 1 and the right hippocampi in Experiments 2-4 were processed for whole-cell western blotting. Tissue was sonicated using the Fisherbrand Model 50 Sonic Dismembrator (Fisher) in 10 μl/mg lysis buffer containing 1 mM EGTA, 1 mM EDTA, 20 mM Tris, 1 mM sodium pyrophosphate tetrabasic decahydrate, 4 mM 4-nitrophenyl phosphate disodium salt hexahydrate, 0.1 μM microcystin, and 1% protease inhibitor cocktail (Sigma-Aldrich). Samples were then centrifuged for 15 min at 1000 x g at 4°C. Bradford protein assays were performed to determine the protein concentration of each sample. Each sample diluted 1:1 with Laemmli Sample Buffer (BioRad) mixed with 350 mM DTT, boiled for 10 min, and stored at −80°C until western blotting.

Fifteen micrograms of cytosolic, membrane, and nuclear protein from each sample, or 25 ug of whole-cell protein from each sample, was loaded onto and separated on a 7.5% TGX SDS-PAGE gel at 250 V for 40 minutes. Molecular weight markers (Precision Plus Protein Standards, BioRad) were included with each run. Proteins were transferred from gels to nitrocellulose membranes at 100 V for 30 minutes. Membranes were blocked with 5% bovine serum albumin (BSA) in 1% Tween 20/Tris-buffered saline (TTBS) with gentle mixing at room temperature for 1 hour. After blocking, membranes were incubated with gentle mixing in primary antibody overnight at 4°C in 1% BSA-TTBS. Samples from cytosolic, membrane, and nuclear compartments were incubated with antibodies for phospho-S118 ERα (1:1000, Abcam) and total ERα (1:1000, Santa Cruz). Samples from cytosolic fractions were incubated with antibodies for cytosolic loading control Enolase (1:1000, Santa Cruz). Samples from membrane fractions were incubated with antibodies for membrane loading control ATP1A1 (1:5000, ProteinTech). Samples from nuclear fractions were incubated with antibodies for nuclear loading control histone 3 (H3; 1:1000, Cell Signaling). Whole-cell tissue samples were incubated with antibodies for phospho-MAPK (1:1000, Cell Signaling) and total p42-MAPK (1:1000, Cell Signaling) which recognize both the p44- and p42-MAPK epitopes, phospho-Akt (1:1000, Cell Signaling), total Akt (1:1000, Cell Signaling), total ERα (1:1000, Santa Cruz), aromatase (1:1000, BioRad), or loading control GAPDH (1:1000, Santa Cruz). Following primary antibody incubation, blots were washed three times for 15 minutes with TTBS. Blots were then incubated with secondary antibodies conjugated to fluorophores in 5% BSA-TTBS for one hour at room temperature with gentle mixing. Secondary antibodies used were StarBright B520 Rabbit (BioRad; 1:5000 for p-S118 ERα, enolase, ATP1A1, H3, total MAPK, total Akt) and StarBright B700 Mouse (BioRad; 1:5000 for total ERα, phosphor-MAPK, phospho-Akt, aromatase, GAPDH). Blots were washed three times for 15 minutes with TTBS, and then imaged on the ChemiDocMP set to channels for StarBright B520 and StarBright B700. MCID Core imaging software was used to quantify optical density for bands of interest.

### Hippocampal estradiol detection

Left hippocampi from Experiments 3-4 were processed for estradiol extraction and measurement via UPLC-MS/MS as previously described and validated (Li et al 2016). This method has recently been shown to sensitively detect estradiol levels in hippocampal tissue from ovariectomized rats treated with estradiol in a dose-dependent manner (Li and Gibbs, 2019). Briefly, tissues were homogenized in a potassium phosphate buffer (0.12M, pH 7.4; 100 mg tissue/mL) containing 4.0 mM MgCl_2_, 4.0 mM Tris and 50 mM sucrose. Samples were spiked with deuterated 17 beta-estradiol and then extracted with n-butyl chloride (Sigma-Aldrich, Inc). The organic layer was dried under nitrogen, then resuspended and derivatized with dansyl chloride in a 1:1 mix of acetonitrile:water (pH 10.5, Sigma-Aldrich, Inc). Samples were then centrifuged and the supernatant transferred to glass vials for UPLC-MS/MS analysis. Calibration curves were prepared in a matrix of 0.2% 2-hydroxypropyl-ß-cyclodextrin (HPCD) and processed the same as the tissue extracts.

Estradiol was eluted using a Waters Acquity UPLC BEH C18, 1.7 µm, 21 x 150 mm reversed-phase column, with an acetonitrile:water (0.1% formic acid) gradient. Detection was in the positive mode. Transitions used for analysis were 506->171 for estradiol, and 511->171 for the internal standard. Note that this method is able to distinguish between 17-alpha and 17-beta estradiol based on retention time. Limit of detectability is 0.009 pmol/mL (2.5 pg/mL) with intra-day and inter-day relative standard deviations of less than 15% at all concentrations.

### Experimental Design and Statistical Analyses

#### Experiment 1

Middle-aged rats were ovariectomized and 10 days later treated them with an acute intracerebroventricular (icv) infusion of either vehicle, IGF-1, or IGF-1 plus letrozole, an aromatase inhibitor that blocks estrogen synthesis. After one (Vehicle, n=9; IGF-1, n=9; IGF-1 + Letrozole, n=10) or 24 (Vehicle, n=10; IGF-1, n=9; IGF-1 + Letrozole, n=9) hours, animals were euthanized and hippocampi were dissected. Right hippocampal tissue was collected and processed for subcellular fractionation and western blotting for phospho-S118 ERα and total ERα. Left hippocampal tissue was collected and processed for whole-cell western blotting for phospho-p42-MAPK/total p42-MAPK and phospho-Akt/total Akt to determine if blocking neuroestrogen synthesis decreases MAPK and PI3K-Akt signaling in animals simultaneously treated with IGF-1.

#### Experiment 2

Middle-aged rats were trained on the 8-arm radial maze for 21 days before undergoing ovariectomy. Ten days after ovariectomy, rats were implanted with a cannula and mini-pump which chronically delivered either vehicle (n=11), IGF-1 (n=9), or IGF-1 plus letrozole (n=11) to the lateral ventricle for the duration of the experiment. Animals were then tested on delay trials (1 min, 1hr, 3 hr, 4 hr, 5 hr) in the 8-arm radial maze to test hippocampal-dependent spatial memory. Two days after the final day of delay testing, animals were euthanized, and right hippocampal tissue was collected and processed for whole-cell western blotting for phospho-p42-MAPK/total p42-MAPK and phospho-Akt/total to determine if chronic letrozole treatment impacts hippocampal activation of the MAPK and PI3K-AkT pathways in animals treated with IGF-1.

#### Experiment 3

Middle-aged rats were ovariectomized and immediately implanted with subcutaneous vehicle capsules for forty days (to match subsequent estradiol treatments in Experiment 4). Forty days later, capsules were removed. Animals were allowed to age for sixty more days following capsule removal before behavioral training begin, resulting in a total of one-hundred days between removal of estrogens (ovariectomy) and behavioral training. Following that sixty-day waiting period, animals were trained on the radial-arm maze for 24 days. Animals then underwent stereotaxic surgery and were implanted with a cannula and mini-pump that chronically delivered either vehicle (n=10), IGF-1 receptor antagonist JB1 (n=10), aromatase inhibitor letrozole (n=9), or JB1 and letrozole (n=9). Animals were tested on delay trials (No delay, 1 min, 1 hr) in the radial-arm maze to test hippocampal-dependent spatial memory. Two days after the final day of delay testing, animals were euthanized and right hippocampal tissue was collected and processed for western blotting for ERα, aromatase, phospho-p42-MAPK, total p42-MAPK, phospho-Akt, and total-Akt were performed. Left hippocampal tissues were collected and processed for estradiol detection via UPLC-MS/MS.

#### Experiment 4

Middle-aged rats were ovariectomized and immediately implanted with subcutaneous estradiol capsules. Forty days later, capsules were removed. Animals were allowed to age for one-hundred more days following capsule removal before behavioral training begin, resulting in a total of one-hundred days between removal of estrogens (capsule removal) and behavioral training. Following the one-hundred day waiting period, animals were trained on the radial arm maze for 24 days. Animals then underwent stereotaxic surgery and were implanted with a cannula and mini-pump that chronically delivered either vehicle (n=9), IGF-1 receptor antagonist JB1 (n=8), aromatase inhibitor letrozole (n=9), or JB1 and letrozole (n=9). Animals were tested on delay trials (No delay, 1 min, 1 hr, 2hr, 3hr) in the 8-arm radial maze to test hippocampal-dependent spatial memory. Two days after the final day of delay testing, animals were euthanized and right hippocampal tissue was collected and processed for western blotting for ERα, aromatase, phospho-p42-MAPK, total p42-MAPK, phospho-Akt, and total-Akt were performed. Left hippocampal tissue were collected and processed for estradiol detection via UPLC-MS/MS.

#### Statistical Analyses

All data analyses were performed using SPSS software. Behavioral data were analyzed by Mixed-Design ANOVA comparing Errors of 8 between treatment groups and across delay trials. Subsequent post-hoc testing, as described below, was used as appropriate for between-subject effects. Western blotting and mass spec data were analyzed by One-Way ANOVA comparing optical density and estradiol levels in fmol/mL, respectively, between experiment groups with subsequent post-hoc testing as appropriate.

For experiments with only three experimental groups (Experiment 1 and 2), LSD post-hoc testing was used as appropriate for between-group effects. For experiments with more than three experimental groups (Experiments 3 and 4), a significant main effect of treatment was probed by the Dunnett’s 2-sided post hoc test, which compares treatments with a single control group (Vehicle group). Western data were analyzed by One-Way ANOVA comparing optical density between treatment group and subsequent post-hoc testing as appropriate. For quantification of estradiol levels, two samples from the Letrozole group in Experiment 3 were used for spike and recovery tests to optimize procedures for these set of samples and were therefore excluded from statistical analysis. Additionally, due to the high sensitivity of mass spec detection, extreme statistical outliers as identified by SPSS software (defined as ± 3 × interquartile range from the first or third quartiles for each group) were presumed to indicate sample contamination and therefore were excluded from statistical analyses. Researchers were blind to treatment group during behavioral testing, western blotting, mass spec, and data analysis.

## Results

### Experiment 1: Neuroestrogen synthesis is necessary for IGF-1-mediated phosphorylation and subsequent increase in protein levels of ERα in the hippocampus of recently ovariectomized middle-aged rats

In the absence of ovarian estrogens, IGF-1 activates ERα via ligand-independent mechanisms (Kato et al. 1995; Grissom and Daniel 2016). IGF-1 activation of ERα *in vitro* requires concomitant neuroestrogen synthesis (Pollard and Daniel 2019). The goal of this experiment was to test the hypothesis that neuroestrogen synthesis is required for IGF-1 activation of ERα protein *in vivo*. Recently ovariectomized middle-aged received icv infusions of either vehicle (Veh group), IGF-1 (IGF-1 group), or IGF-1 plus letrozole (IGF-1 + Let group). Total and phosphorylated levels of ERα and the IGF-1 regulated signaling proteins MAPK and AKT, were measured either 1- or 24-hours post-infusion.

*IGF-1-mediated phosphorylation of ERα in cytosolic compartment and subsequent increase in total levels of ERα in the nuclear compartment require neuroestrogen synthesis*.

As a nuclear steroid receptor with the ability to be inserted into cell membranes, the subcellular localization of ERα impacts the receptor’s function. Therefore, we measured levels of phosphorylated and total ERα levels in the cytosol, membrane, and nuclear compartments of hippocampal cells at each time point.

In the cytosolic compartment (Figure 1), there was a main effect of treatment on levels of pS118-ERα (F(2,27)=4.973; p=0.015) at 1 hour after treatment (Figure 1A), with levels of pS118-ERα significantly increased in the IGF-1 treatment group as compared to the vehicle group (p=0.007) and the IGF-1+Let treatment group (p=0.020). This observed increase in phosphorylated ERα in the cytosol after 1 hour is consistent with earlier work in cell cultures demonstrating that peak dimerization (and presumably therefore, nuclear translocation) of ERα does not occur until two hours after estrogen treatment (Powell and Xu, 2008). However, there was no significant difference in cytosolic pS118-ERα levels between the IGF-1+Let and vehicle groups (p=0.611). There was no effect of treatment on total levels of cytosolic ERα 1-hour post-infusion (Figure 1B; F(2,27)=0.553, p=0.582). At the 24-hour time point, there were no effects of treatment on cytosolic levels of pS118-ERα (Figure 1C; F(2,27)=0.292, p=0.750) or total ERα (Figure 1D; F(2,27)=1.408, p=0.263).

**Figure 1.**
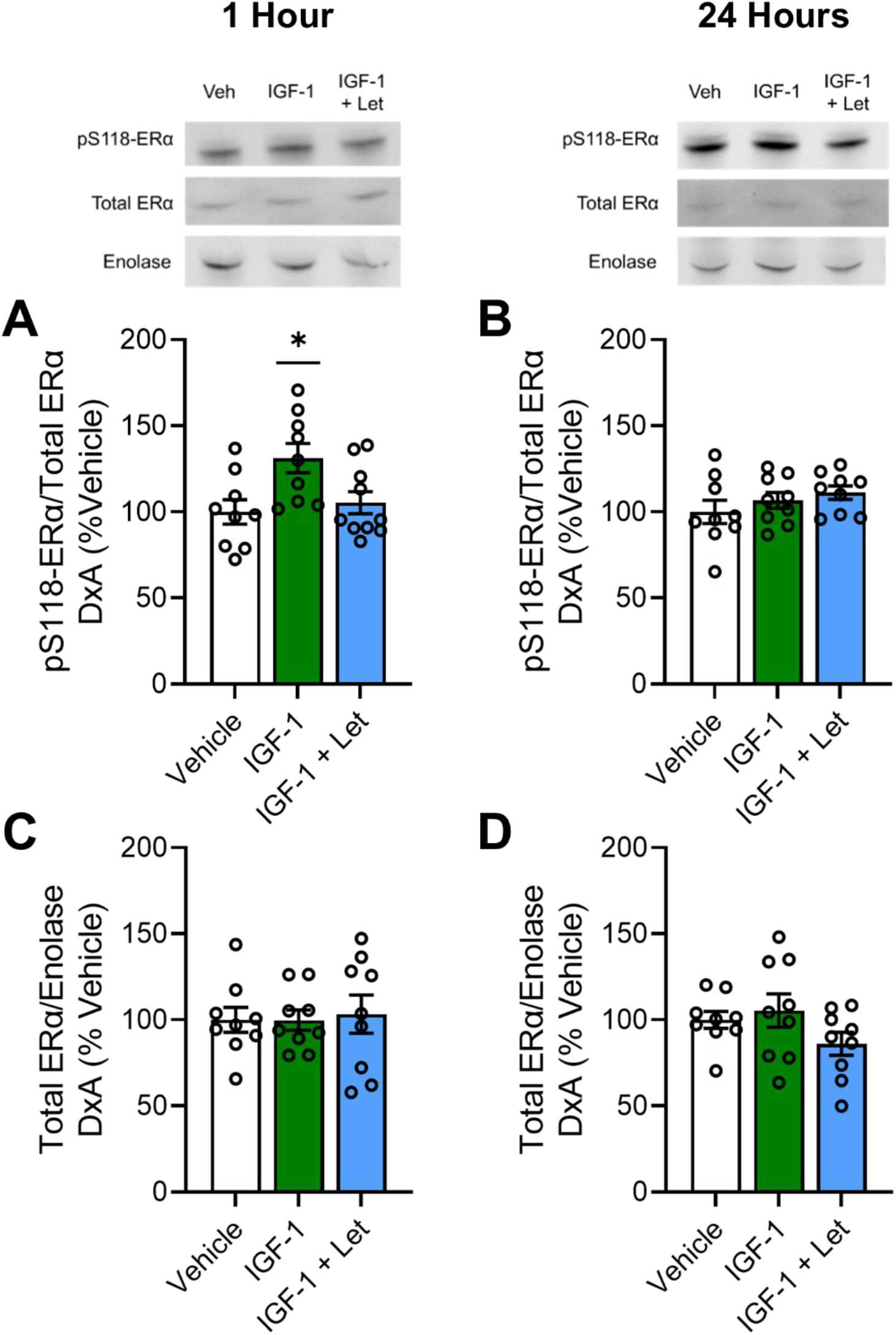
Cytosolic expression of pS118-ERα and total ERα 1-hour or 24-hours post infusion of IGF-1 or IGF-1 + letrozole in the hippocampus of ovariectomized rats. Middle-aged female rats were ovariectomized and given an acute infusion of either vehicle (Veh), insulin-like growth factor-1 (IGF-1), or IGF-1 and the aromatase inhibitor letrozole (IGF-1 + Let) to the lateral ventricle. Either 1 or 24-hours later, hippocampi were processed for subcellular fractionation and western blotting for phosphorylated levels of ERα at S118 (pS118-ERα), total ERα, and cytosolic loading control enolase in the cytosolic fraction of all samples. Levels of pS118 were normalized to total ERα levels % vehicle group, and levels of total ERα were normalized to enolase % vehicle group. A) There was an effect of treatment (*p*<0.05) on pS118-ERα levels in the cytosol compartment 1-hour post infusion, with post hoc testing revealing increased levels in the IGF-1 group as compared to vehicle group. B-D) There was no effect of treatment on pS118-ERα levels 24-hours post infusion (B), nor was there an effect of treatment on total ERα levels either 1-hour (C) or 24-hours (D) post infusion. *p<0.05 vs. Veh

In the membrane compartment (Figure 2), there were no effects of treatment 1-hour post-infusion on levels of pS118-ERα (Figure 2A, F(2,25)=1.243; p=0.307) or total ERα (Figure 2B, F(2,27)=0.875; p=0.429). There were also no effects of treatment 24-hours post-infusion on membrane levels pS118-ERα (Figure 2C, F(2,27)=2.122; p=0.141) or total ERα (Figure 2D, F(2,27)=0.528; p=0.596).

**Figure 2.**
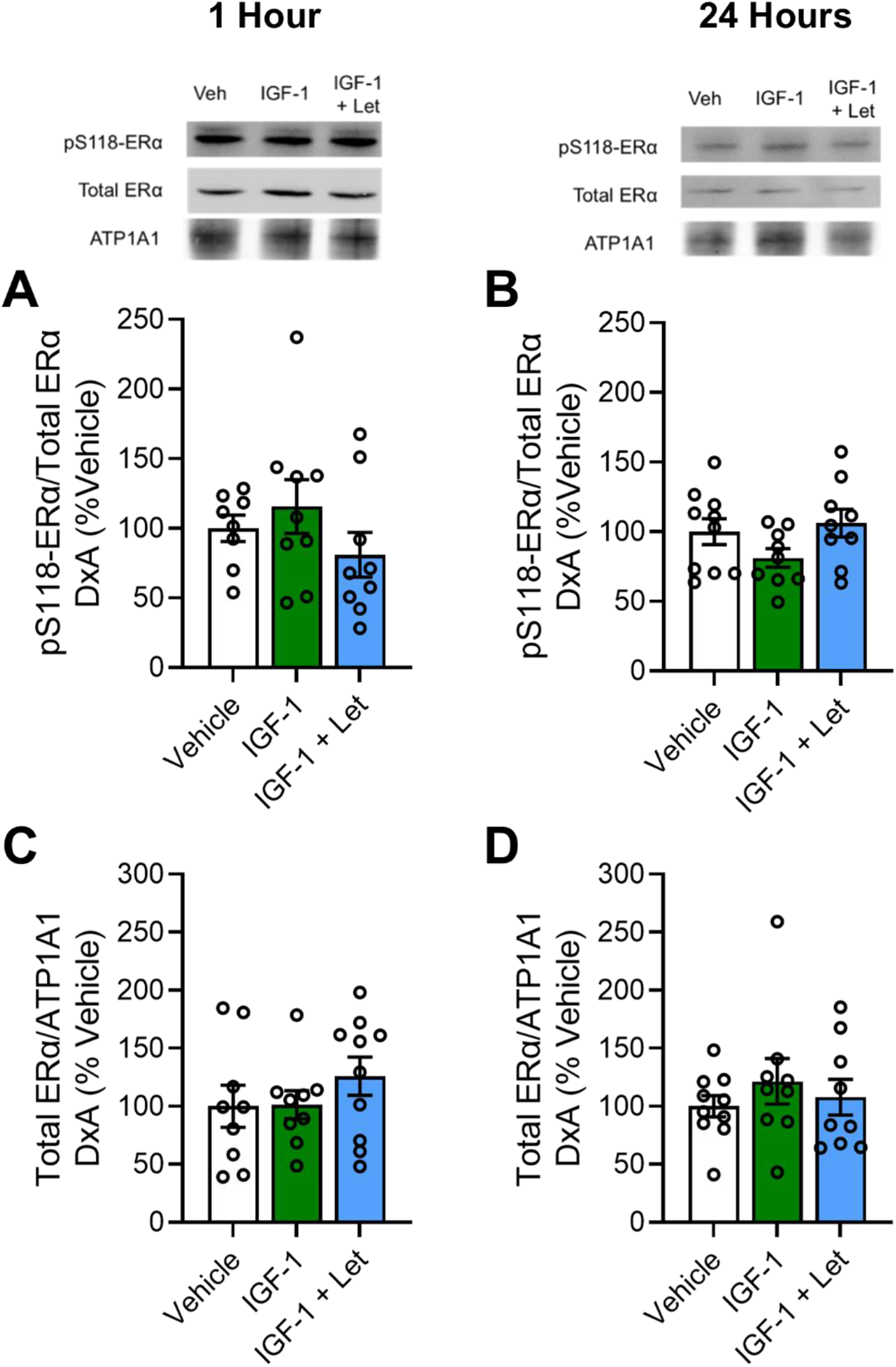
Membrane expression of pS118-ERα and total ERα 1-hour or 24-hours post infusion of IGF-1 or IGF-1 + letrozole in the hippocampus of ovariectomized rats. Middle-aged female rats were ovariectomized and given an acute infusion of either vehicle (Veh), insulin-like growth factor-1 (IGF-1), or IGF-1 and the aromatase inhibitor letrozole (IGF-1 + Let) to the lateral ventricle. Either 1 or 24-hours later, hippocampi were processed for subcellular fractionation and western blotting for phosphorylated levels of ERα at S118 (pS118-ERα), total ERα, and membrane loading control ATP1A1 in the membrane fraction of all samples. Levels of pS118-ERα were normalized to total ERα levels % vehicle group, and levels of total ERα were normalized to ATP1A1 % vehicle group. A-B) There was no effect of treatment on membrane pS118-ERα levels either 1-hour (A) or 24-hours (B) post infusion. C-D) There was no effect of treatment on membrane total ERα levels either 1-hour (C) or 24-hours (D) post infusion.

In the nuclear compartment (Figure 3), there were no effects of treatment on levels of pS118-ERα (Figure 3A; F(2,27)=0.095, p=0.910) or total ERα (Figure 3B; F(2,27)=0.202, p=0.818) 1-hour post-infusion. At the 24-hour timepoint, there was no effect of treatment on pS118-ERα levels in the nuclear compartment (Figure 3C; F(2,27)=0.084, p=0.919). However, there was an effect of treatment on total ERα levels in the nuclear compartment (Figure 3D; F(2,27)=3.915, p=0.033) 24-hours post-infusion, with the IGF-1 treatment group showing significantly higher levels as compared to the vehicle group (p=0.011) and near significantly higher levels as compared to the IGF-1+Let group (p=0.079). There was no significant difference in total ERα levels between the vehicle and IGF-1+Let treatment groups (p=0.388).

**Figure 3.**
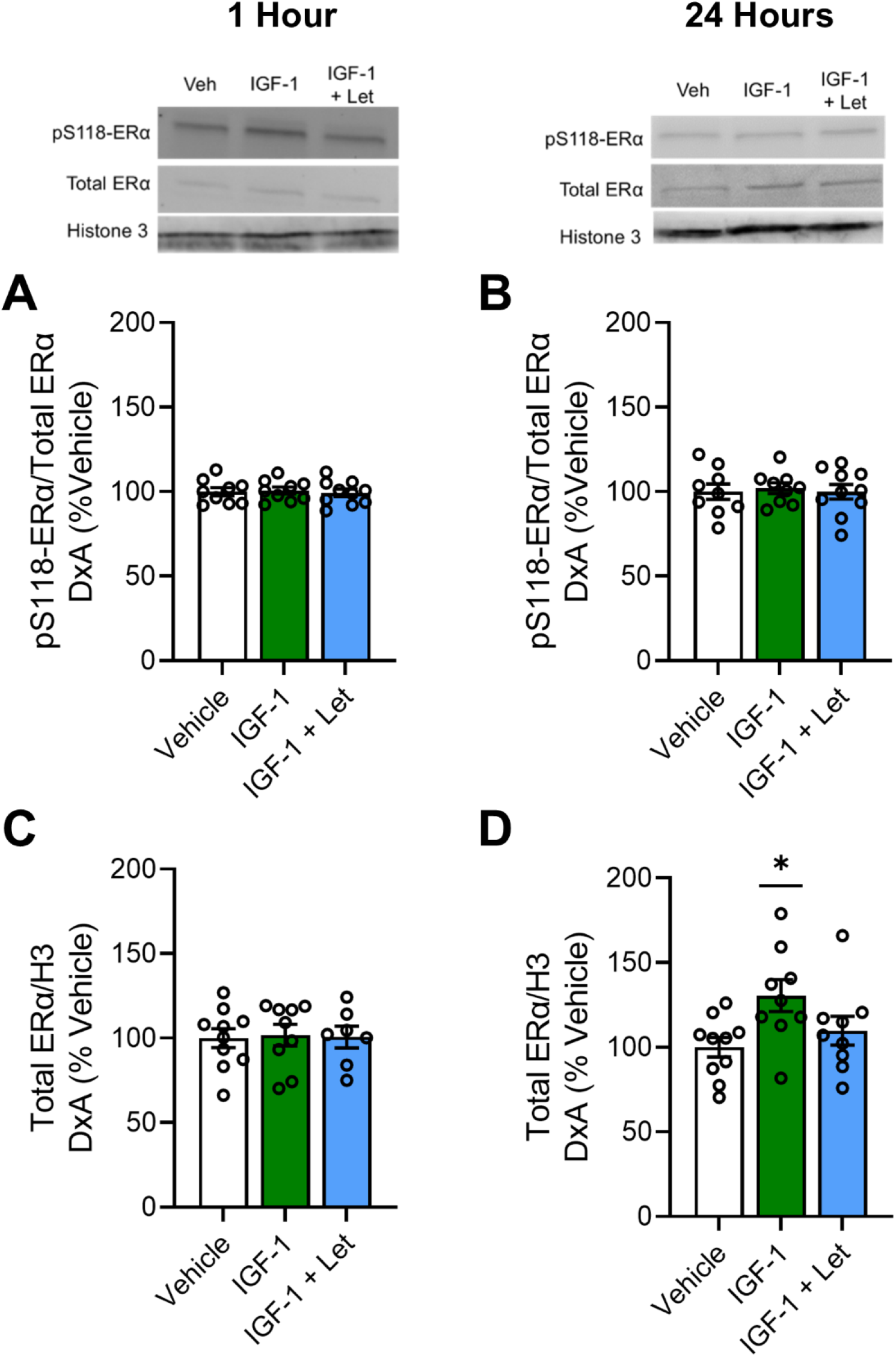
Nuclear expression of pS118-ERα and total ERα 1-hour or 24-hours post infusion of IGF-1 or IGF-1 + letrozole in the hippocampus of ovariectomized rats. Middle-aged female rats were ovariectomized and given an acute infusion of either vehicle (Veh), insulin-like growth factor-1 (IGF-1), or IGF-1 and the aromatase inhibitor letrozole (IGF-1 + Let) to the lateral ventricle. Either 1 or 24-hours later, hippocampi were processed for subcellular fractionation and western blotting for phosphorylated levels of ERα at S118 (pS118-ERα), total ERα, and nuclear loading control CREB in the nuclear fraction of all samples. Levels of pS118 were normalized to total ERα levels % vehicle group, and levels of total ERα were normalized to CREB % vehicle group. A-B) There was no effect of treatment on pS118-ERα levels in the nuclear compartment either 1-hour (A) or 24-hours (B) post infusion. C) There was no effect of treatment on pS118-ERα levels 24-hours post infusion. D) There was an effect of treatment (*p*<0.05) on total ERα levels 24-hours post infusion, with post hoc testing revealing increased levels in the IGF-1 group as compared to the Veh group. *p<0.05 vs. Veh

*In parallel to impacts of IGF-1 on ERα activation, IGF-1-mediated activation of MAPK, but not Akt requires neuroestrogen synthesis*.

In order to determine if IGF-1 activation of ERα occurs via the MAPK or PI3K-Akt signaling pathways and if IGF-1 activation of pathways require neuroestrogen synthesis, we measured phosphorylated and total levels of p44-MAPK (ERK-1), p42-MAPK (ERK-2), and Akt.

One hour post infusion, there was a main effect of treatment on phosphorylation of both p44 (Figure 4A; F(2,27)= 35.750, p=4.65×10-8) and p42 (Figure 4B; F(2,27)= 21.720, p=3.41×10-6) phospho-sites of MAPK. Post hoc testing revealed a significant increase in phosphorylation of both phospho-sites of MAPK in the IGF-1 treatment group as compared to the Vehicle group (p44-MAPK, p=2.39×10-7; p42-MAPK, p=2.59×10-6) and the IGF-1 + Let group (p44-MAPK, p=5.20×10-8; p42-MAPK, p=1.48×10-5). There was no difference between the IGF-1 + Let group and the Veh group on phosphorylation of either p44 (p=0.647) or p42 (p=0.407) MAPK levels.

**Figure 4.**
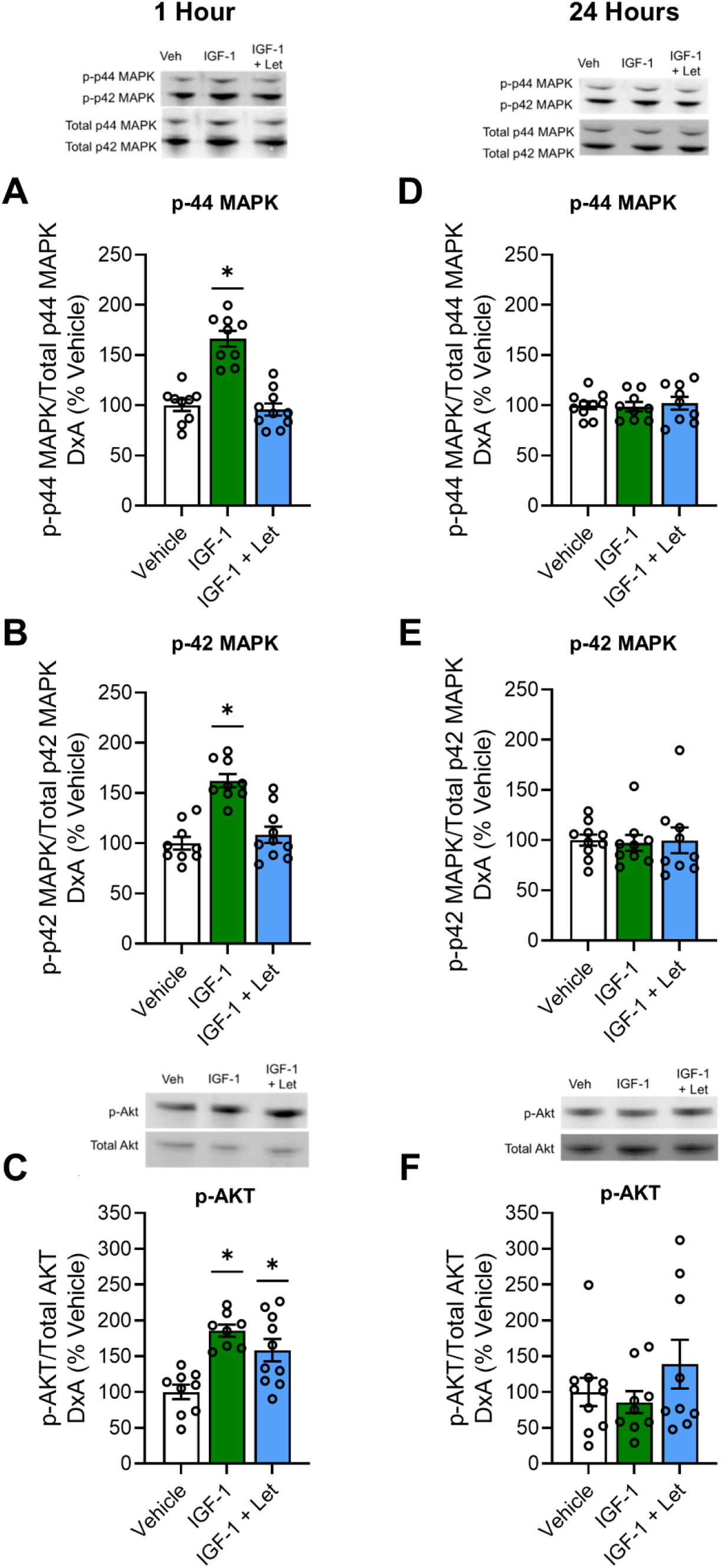
Hippocampal MAPK and Akt pathway activation 1-hour or 24-hours post infusion of IGF-1 or IGF-1 + letrozole. Middle-aged female rats were ovariectomized and given an acute infusion of either vehicle (Veh), insulin-like growth factor-1 (IGF-1), or IGF-1 and the aromatase inhibitor letrozole (IGF-1 + Let) to the lateral ventricle. Either 1 or 24-hours later, hippocampi were processed for western blotting for phosphorylated and total levels of p44-MAPK, p42-MAPK, and Akt. Phosphorylated levels were normalized to the total protein levels and expressed as a percentage of the Veh group mean. A-D) One hour post infusion, there was a main effect of treatment (*p*<0.05) on phosphorylated levels of p44-MAPK (A) and p42-MAPK (B), with post hoc testing revealing increased phosphorylation of both MAPK phosphorylation sites in the IGF-1 group as compared to the Veh group. There was no effect of treatment on phosphorylated levels of p44-MAPK (D) or p42-MAPK (E) 24-hours post infusion. C) One hour post infusion, there was an effect (*p<*0.05) of treatment on phosphorylated levels of Akt, with post hoc testing revealing increased levels in both the IGF-1 and IGF-1 + Let groups as compared to the Veh group. F). There was no effect of treatment on phosphorylated levels of Akt 24-hours post infusion. *p<0.05 vs. Veh

One hour post infusion, there was a main effect of treatment on phosphorylation of Akt (Figure 4C; F(2,27)= 7.552, p=0.003). Post hoc testing revealed a significant increase in phosphorylation of Akt in the IGF-1 group (p=0.001) and the IGF-1 + Let group (p=0.006) as compared to the Veh group.

Twenty-four hours post infusion, there was no main effect of treatment on phosphorylation of p44 MAPK (Figure 4D; F(2,27)= 0.120, p=0.888), p42 MAPK (Figure 4E; F(2,27)= 0.031, p=0.970), or Akt (Figure 4F; F(2,27)= 1.247, p=0.297).

### Experiment 2: Neuroestrogen synthesis is necessary for IGF-1-mediated enhancement of spatial memory in recently ovariectomized middle-aged rats

Experiment 1 revealed that IGF-1 activation of the MAPK signaling pathway, associated phosphorylation of ERα, and subsequent increase in levels of ERα in the hippocampus were blocked by letrozole, suggesting that they require neuroestrogen synthesis. In Experiment 2, we determined the functional consequences of these effects by testing the hypothesis that the ability of IGF-1 to impact hippocampal-dependent memory in recently ovariectomized rats also requires concomitant neuroestrogen synthesis. Additionally, we determined if IGF-1 activation of signaling pathways in these behaviorally tested animals would parallel effects on memory. Following training on the radial maze, recently ovariectomized middle-aged rats received chronic icv treatment of either vehicle (Veh group), IGF-1 (IGF-1 group), or IGF-1 plus letrozole (IGF-1 + Let group) and were tested on delay trials in the maze. Following maze testing, hippocampal levels of MAPK and PI3K-Akt pathway activation were measured.

*IGF-1 enhances memory performance of recently ovariectomized rats on the radial-arm maze, an enhancement that requires neuroestrogen synthesis*.

Following recovery from stereotaxic surgeries, animals were tested across multiple increasing delays (1 hr, 3 hr, 4 hr, 5 hr) on the 8-arm radial maze test. As illustrated in Figure 5, Mixed-Design ANOVA revealed a main effect of delay (F(3,84)=4.257; p=0.008) and a main effect of treatment (F(2,28)=5.245; p=0.012) on radial-arm maze performance. Post hoc testing revealed significantly fewer errors of 8 across delays in the IGF-1 group as compared to both the Veh group (p=0.004) and the IGF-1 + Let group (p=0.029). There was no difference between the Veh group and the IGF-1 + Let group. There was no significant interaction between delay and treatment (F(6,84)=1.555; p=0.171).

**Figure 5.**
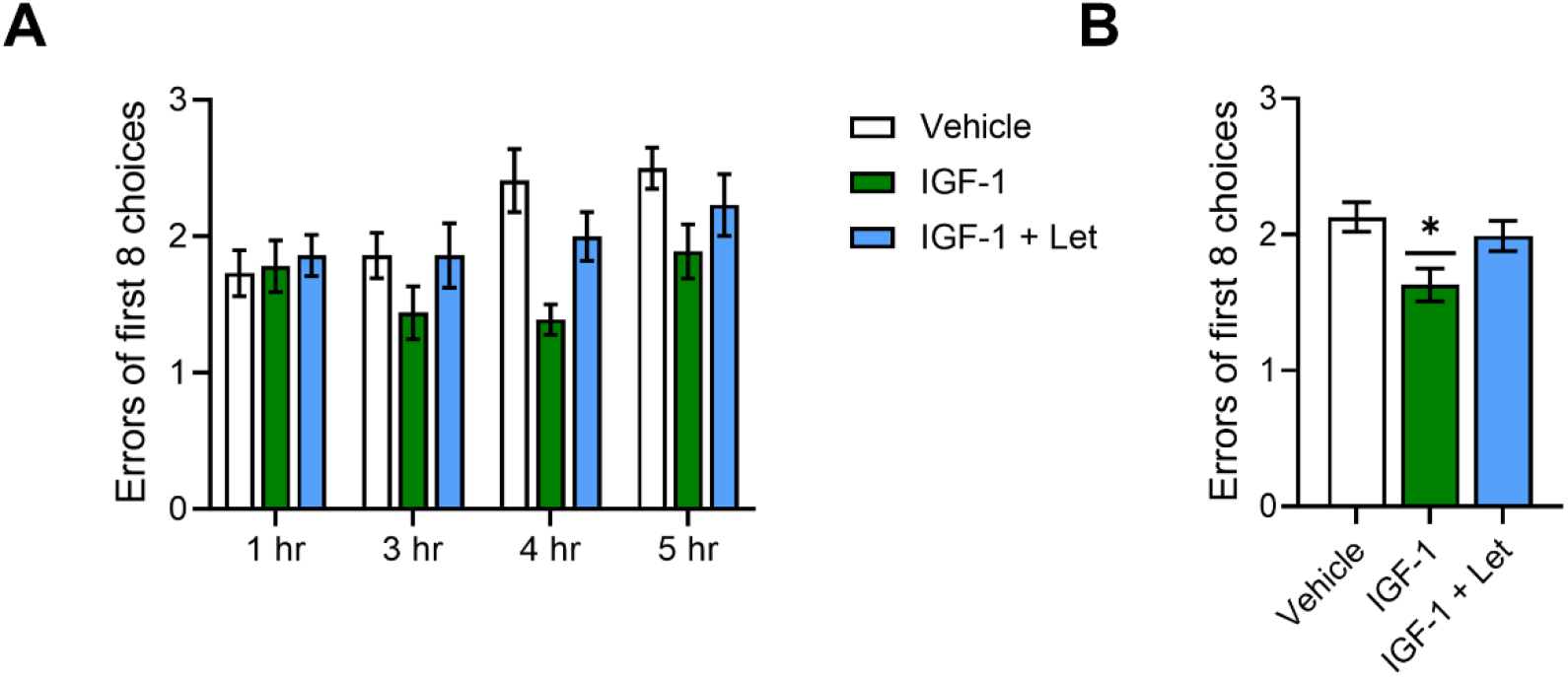
Impacts of chronic IGF-1 or IGF-1 + Letrozole treatment on performance on the hippocampal-dependent radial-arm maze task. Middle-aged female rats were trained on the 8-arm radial maze task and subsequently ovariectomized and treated with either vehicle (Veh), insulin-like growth factor-1 (IGF-1) or IGF-1 plus the aromatase inhibitor letrozole (IGF-1+Let) and tested on the maze using delays of 1, 3, 4, and 5 hours. Data represent the number of incorrect choices made in the first eight choices averaged across two days of testing at each delay. A) There was a main effect of delay (*p*<0.05) on performance across groups, with post hoc testing revealing errors increased as delays became longer. There was no significant interaction between delay x treatment. B) There was a main effect of treatment (*p*<0.05) on performance averaged across all delays, with post hoc testing revealing significantly fewer errors in the IGF-1 group as compared to the Veh group. *p<0.05 vs. Veh

*IGF-1 activation of MAPK, but not Akt signaling requires neuroestrogen synthesis in recently ovariectomized, behaviorally tested rats*

Following chronic icv treatment with either IGF-1 or IGF-1 + Let, there was a main effect of treatment on phosphorylation of p44-MAPK (Figure 6A; F(2,30)= 32.346, p=5.27×10-7) and a near significant effect of treatment on phosphorylation of p42-MAPK (Figure 6B; F(2,30)= 3.106, p=0.060). Post hoc testing revealed a significant increase in phosphorylation of p44-MAPK levels in the IGF-1 group as compared to both the Veh group (p=1.28×10-7) and the IGF-1 + Let group (p=8.74×10-8). There was also a significant increase in phosphorylation of p42-MAPK levels in the IGF-1 group as compared to the Veh group (p=0.020) and a statistically trending increase as compared to the IGF-1 + Let group (p=0.104). There was no difference between the IGF-1 + Let group and the Veh group on phosphorylation of either p44 (p=0.878) or p42 (p=0.418) MAPK levels.

**Figure 6.**
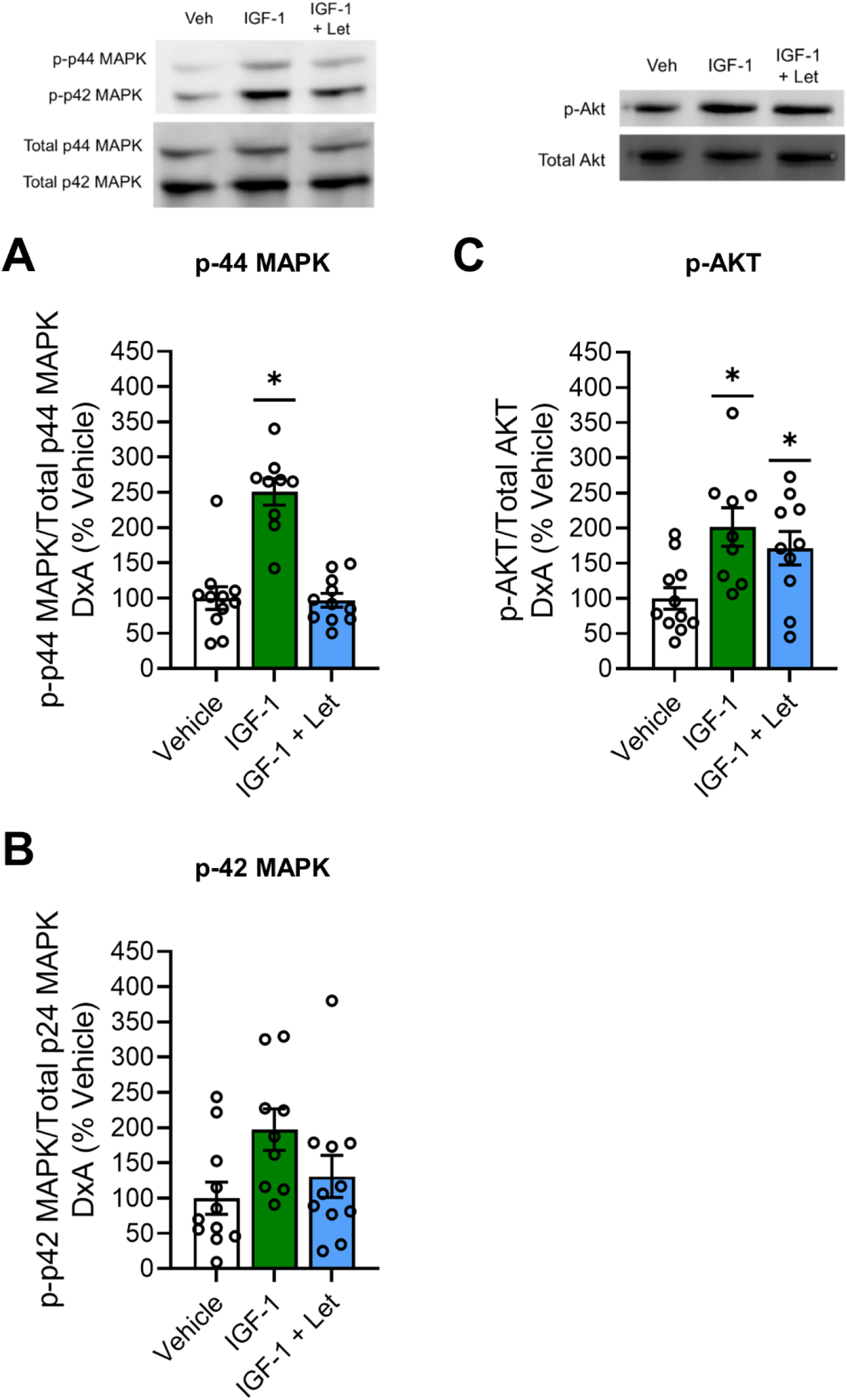
Impacts of chronic IGF-1 or IGF-1 + Letrozole treatment on hippocampal MAPK and Akt pathway activation. Middle-aged female rats were ovariectomized and treated with either vehicle (Veh), insulin-like growth factor-1 (IGF-1) or IGF-1 plus the aromatase inhibitor letrozole (IGF-1+Let). After rats were tested on the radial-arm maze, hippocampi were dissected and processed for western blotting for phosphorylated and total levels of p44-MAPK, p42-MAPK, and Akt. Phosphorylated levels were normalized to the total protein levels and expressed as a percentage of the Veh group mean. There was an effect of treatment (*p*<0.05) on phosphorylated levels of p44-MAPK (A) and p42-MAPK (B), with post hoc testing revealing increased phosphorylation of both MAPK phosphorylation sites in the IGF-1 group as compared to the Veh group. There was an effect of treatment (*p*<0.05) on phosphorylated levels of Akt (C), with post hoc testing revealing increased phosphorylation in both the IGF-1 and IGF-1+Let groups as compared to Veh group. *p<0.05 vs. Veh

As illustrated in Figure 6C, there was a main effect of treatment on phosphorylation of Akt (F(2,30)= 4.803, p=0.016). Post hoc testing revealed a significant increase in phosphorylation of Akt in the IGF-1 group (p=0.012) and the IGF-1 + Let group (p=0.013) as compared to the Veh group. There was no difference in levels of phosphorylation of Akt between the IGF-1 and IGF-1 + Let groups (p=0.907).

### Experiment 3: Long-term ovarian hormone deprivation disrupts interactions between IGF-1 and neuroestrogen signaling resulting in detrimental impact of endogenous IGF-1 receptor activity on memory

Experiments 1 and 2 revealed that IGF-1-enhancement of hippocampal-dependent memory and elevation of phosphorylated and total hippocampal levels of ERα in recently ovariectomized rats requires concomitant neuroestrogen synthesis. Furthermore, results suggest that these effects are mediated via activation of the MAPK and not the PI3K-Akt signaling pathway. Previous evidence indicates that neuroestrogen synthesis is regulated by circulating estrogens (Nelson et al 2016) and therefore, not surprisingly, long-term ovariectomy results in decreased aromatase expression and neuroestrogen levels (Chen et al. 2021; Ma et al 2020). In Experiment 3, we aimed to determine the implications of decreased level of neuroestrogens resulting from long-term ovariectomy on IGF-1 signaling effects in the hippocampus. We employed a rat model of menopause in which no post-ovariectomy estradiol was administered, modeling women who do not use menopausal hormone therapy. Because we aimed to assess effects of long-term loss of ovarian function on endogenous IGF-1 signaling and subsequent impact for cognitive aging, we chose to antagonize IGF-1 here rather than exogenously administer IGF-1 as was done in Experiments 1 and 2. Long-term ovariectomized rats (100 days) that received no estradiol treatment received chronic icv delivery of vehicle (Veh group) the IGF-1R antagonist JB1 (JB1 group), aromatase inhibitor letrozole (Let group), or JB1 plus letrozole (JB1+Let group) and were tested on the radial-maze task. Hippocampal levels of MAPK and PI3K-Akt pathway activation were measured. Finally, hippocampal expression of ERα, aromatase—the enzyme that converts testosterone to estradiol—and estradiol levels were measured.

*Antagonizing IGF-1 receptor activity unexpectedly enhances spatial memory suggesting that long-term ovariectomy leads to negative impacts of IGF-1 signaling on memory. Inhibition of neuroestrogens reverses the enhancement, but has no impact on its own*.

As illustrated in Figure 7, Mixed-Design ANOVA revealed a main effect of delay (F(2,68)=6.713; p=0.002) and a main effect of treatment (F(3,34)=3.702; p=0.021) on radial-arm maze performance. Post hoc testing revealed significantly fewer errors of 8 across delays in the JB1 group as compared to the Veh group (p=0.012). There was no difference between the Veh group and the Let group (p=0.417) or between the Veh group and the JB1 + Let group (p=0.978). There was no significant interaction between delay and treatment (F(6,68)=0.419; p=0.864).

**Figure 7.**
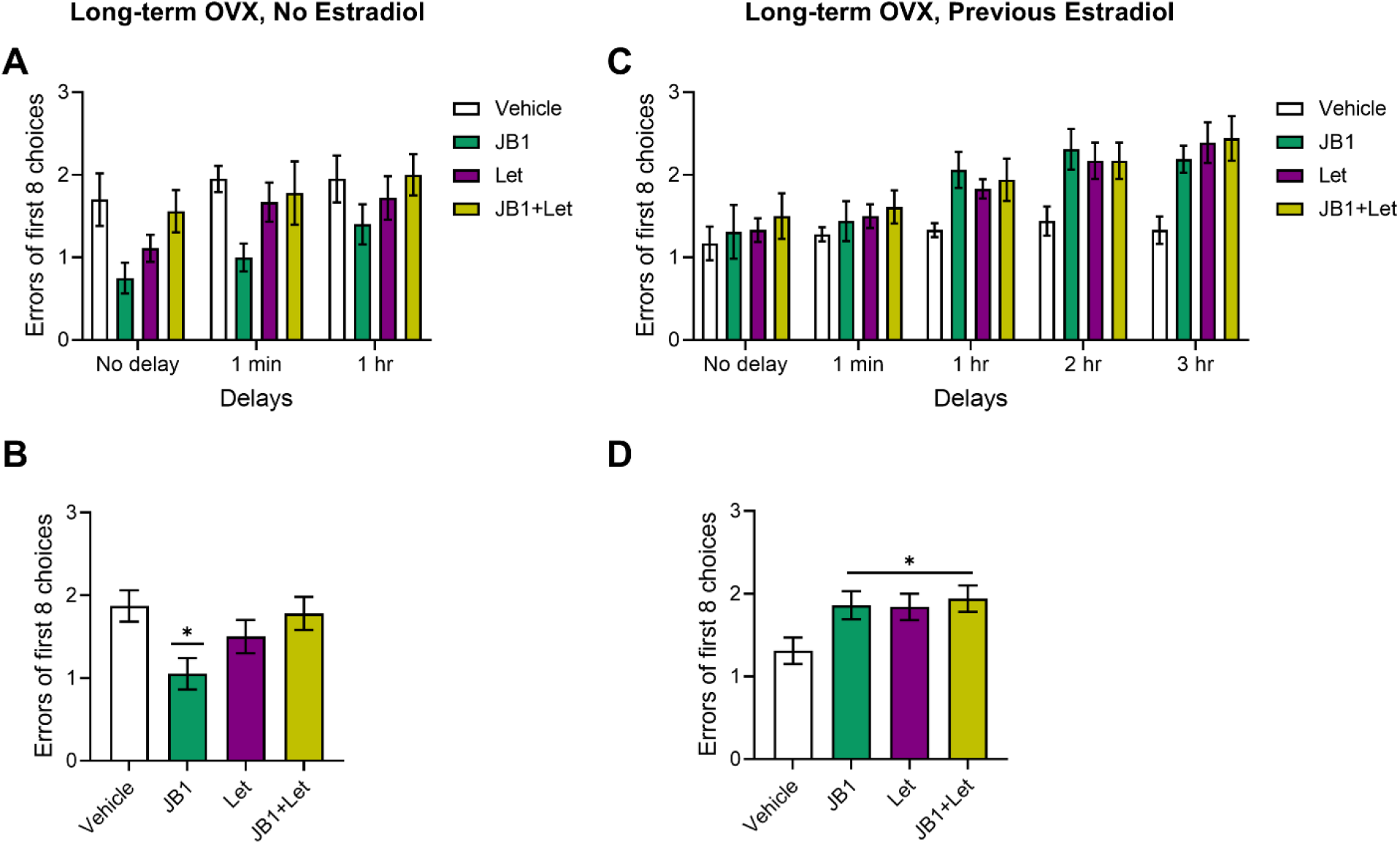
Impacts of chronic JB1, Letrozole, or JB1+Letrozole treatment on radial-arm maze performance in long-term ovariectomized rats with or without previous midlife estradiol exposure. Middle-aged female rats were ovariectomized and immediately implanted with vehicle (Long-term OVX, No Estradiol) or estradiol (Long-term OVX, Previous Estradiol) capsules. Forty days later, capsules were removed. One-hundred days following loss of circulating estrogens (either via ovariectomy in Long-term OVX, No Estradiol group or removal of estradiol capsule in Long-term OVX, Previous Estradiol group), rats were trained on the 8-arm radial maze task and Following training, rats were treated with chronic i.c.v. administration of either vehicle (Veh), the IGF-1R antagonist JB1 (JB1), the aromatase inhibitor letrozole (Let), or JB1 and letrozole (JB1+Let) and tested on the maze using delays (Long-term OVX, No Estradiol: No Delay, 1 minute, and 1 hour; Long-term OVX, Previous Estradiol: No Delay, 1 minute, 1 hour, 2 hour, and 3 hour). Data represent the number of incorrect choices made in the first eight choices averaged across two days of testing at each delay. A) Following long-term ovariectomy with no estradiol exposure, there was a main effect of delay (*p*<0.05) on performance across groups, with post hoc testing revealing errors increased as delays became longer. There was no significant interaction between delay x treatment. B) There was a main effect of treatment (*p*<0.05) on performance averaged across all delays following long-term ovariectomy with no estradiol, with post hoc testing revealing significantly fewer errors in the JB1 group as compared to the Veh group. C) Following long term ovariectomy with previous estradiol exposure, there was a main effect of delay (p<0.05) on performance across groups, with post hoc testing revealing errors increased as delays became longer. There was no significant interaction between delay x treatment. D) There was a main effect of treatment (p<0.05) on performance averaged across all delays following previous estradiol exposure, with post hoc testing revealing significantly more errors in the JB1, Let, and JB1+Let groups as compared to the Veh group. *p<0.05 vs. Veh

*Antagonizing IGF-1 receptor activity increases MAPK signaling and decreases Akt signaling suggesting that under conditions of long-term ovarian hormone deprivation, PI3K-Akt signaling pathway predominates. Inhibition of neuroestrogen synthesis reserves JB1-induced effects on MAPK, but not on Akt*.

After chronic treatment with either JB1, letrozole, or JB1 plus letrozole following long-term ovariectomy, there was an effect of treatment on phosphorylation of both p44-MAPK (Figure 8A; F(3,37)= 5.367, p=0.004) and p42-MAPK (Figure 8B; F(3,37)= 10.793, p=3.90×10^-5^). Post hoc testing revealed a significant increase in phosphorylation of p44-MAPK (p=0.015) and p42-MAPK (p=4.34×10^-4^) levels in the JB1 group as compared to the Veh group. There were no differences between the Veh group and the Let group for either p44-MAPK (p=1.00) or p42-MAPK (p=0.995), nor were there any differences between the Veh group and the JB1+Let group for either p44-MAPK (p=0.805) or p42-MAPK (p=0.709).

**Figure 8.**
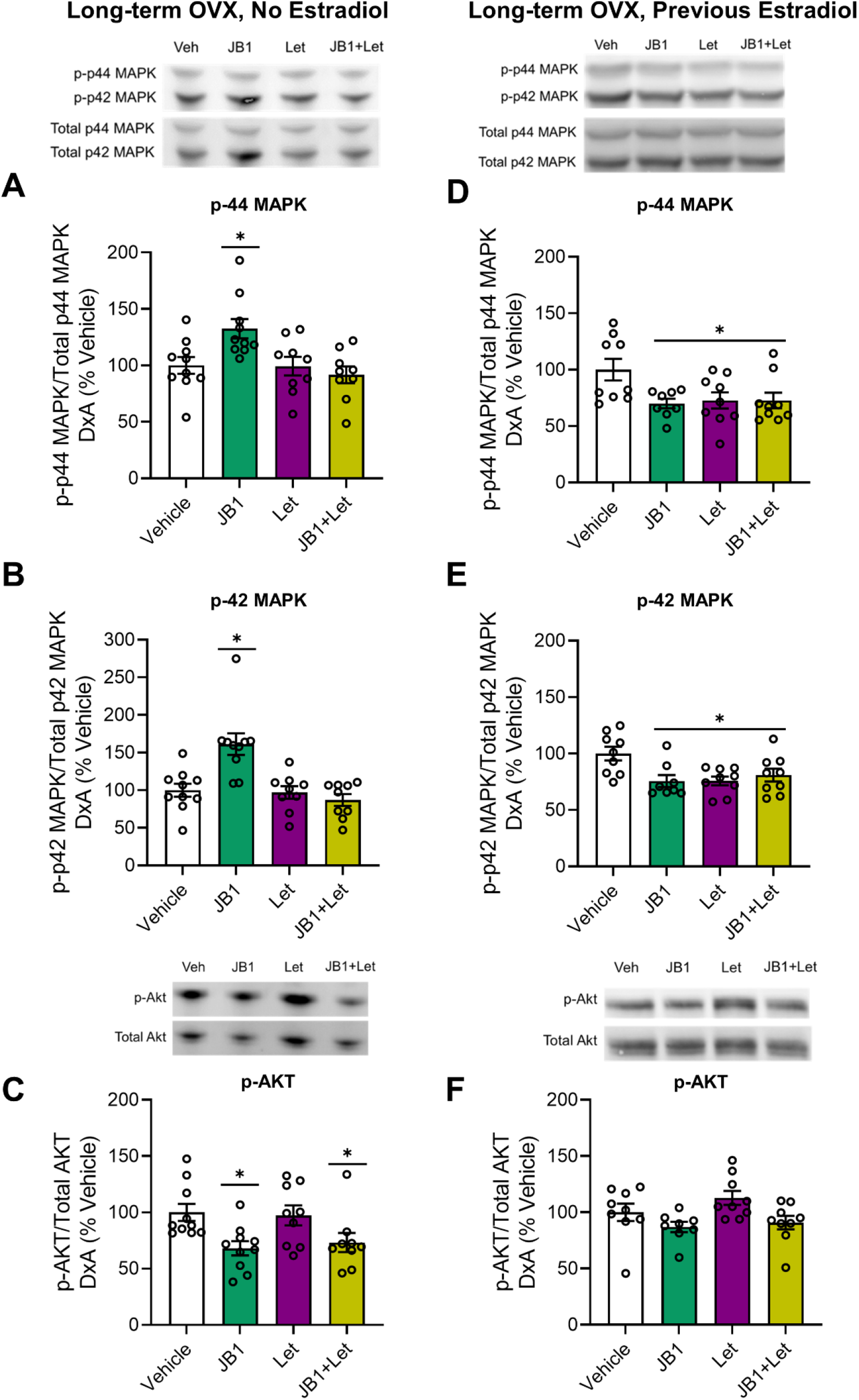
Impacts of chronic JB1, Letrozole, or JB1+Letrozole treatment on hippocampal MAPK and Akt pathway activation in long-term ovariectomized rats with or without previous midlife estradiol exposure. Middle-aged female rats were ovariectomized and immediately implanted with vehicle (Long-term OVX, No Estradiol groups) or estradiol (Long-term OVX, Previous Estradiol groups) capsules. Forty days later, capsules were removed. One-hundred days following loss of circulating estrogens (either via ovariectomy in Long-term OVX, No Estradiol group or removal of estradiol capsule in Long-term OVX, Previous Estradiol group), rats were trained on the 8-arm radial maze task and subsequently treated with chronic i.c.v. administration of either vehicle (Veh), the IGF-1R antagonist JB1 (JB1), the aromatase inhibitor letrozole (Let), or JB1 and letrozole (JB1+Let). After rats were tested on the radial-arm maze, hippocampi were dissected and processed for western blotting for phosphorylated and total levels of p44-MAPK, p42-MAPK, and Akt. Phosphorylated levels were normalized to the total protein levels and expressed as a percentage of the Veh group mean. Following long-term ovariectomy with no estradiol exposure, there was an effect of treatment (*p*<0.05) on phosphorylated levels of p44-MAPK (A) and p42-MAPK (B), with post hoc testing revealing increased phosphorylation of both MAPK phosphorylation sites in the JB1 group as compared to the Veh group. There was also an effect of treatment (*p*<0.05) on phosphorylated levels of Akt (C), with post hoc testing revealing decreased phosphorylation in both the JB1 and JB1+Let groups as compared to Veh group. Following long-term ovariectomy with previous estradiol exposure, there was an effect of treatment (p<0.05) on phosphorylated levels of p44-MAPK (D) and p42-MAPK (E), with post hoc testing revealing decreased phosphorylation of both MAPK phosphorylation sites in the JB1, Let, and JB1+Let groups as compared to the Veh group. There was also an effect of treatment (p<0.05) on phosphorylated levels of Akt (F), with post hoc testing revealing no significant difference between the treatment groups and the Veh group. *p<0.05 vs. Veh

As illustrated in Figure 8C, there was an effect of treatment on phosphorylation of Akt (F(3,37)= 4.437, p=0.010). Post hoc testing revealed a significant decrease in phosphorylation of Akt in the JB1 group (p=0.015) and a near significant decrease in phosphorylation of Akt in the JB1+Let group (p=0.056) as compared to the Veh group. There was no difference in levels of phosphorylation of Akt between the Veh group and the Let group (p=0.990).

*Antagonism of IGF-1 receptors increases protein levels of ERα and aromatase in the hippocampus, effects reversed by inhibition of neuroestrogen synthesis*.

As illustrated in Figure 9A, there was an effect of treatment on hippocampal ERα levels (F(3,37)= 4.202, p=0.012). Post hoc testing revealed a significant increase in ERα expression in the JB1 group (p=0.011) as compared to the Veh group. There was no difference in ERα levels between the Veh group and the Let group (p=0.940) or between the Veh group and the JB1+Let group (p=0.997).

**Figure 9.**
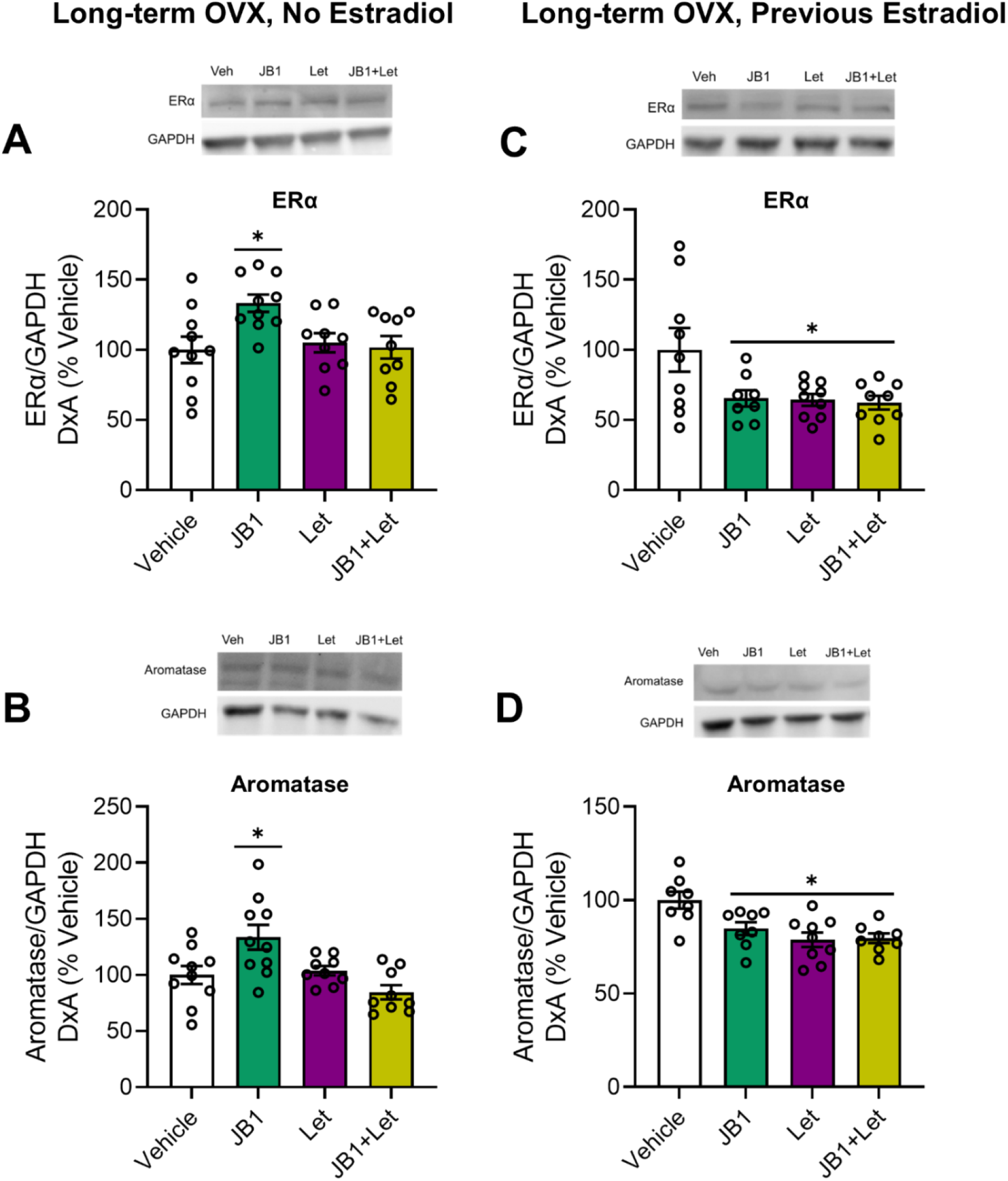
Impacts of chronic JB1, Letrozole, or JB1+Letrozole treatment on hippocampal ERα and aromatase levels in long-term ovariectomized rats with or without previous midlife estradiol exposure. Middle-aged female rats were ovariectomized and immediately implanted with vehicle (Long-term OVX, No Estradiol groups) or estradiol (Long-term OVX, Previous Estradiol groups) capsules. Forty days later, capsules were removed. One-hundred days following loss of circulating estrogens (either via ovariectomy in Long-term OVX, No Estradiol group or removal of estradiol capsule in Long-term OVX, Previous Estradiol group), rats were trained on the 8-arm radial maze task and subsequently treated with chronic i.c.v. administration of either vehicle (Veh), the IGF-1R antagonist JB1 (JB1), the aromatase inhibitor letrozole (Let), or JB1 and letrozole (JB1+Let). After rats were tested on the radial-arm maze, hippocampi were dissected and processed for western blotting for ERα, aromatase, and loading control GAPDH. ERα and aromatase levels were normalized to GAPDH levels and expressed as a percentage of the Veh group mean. Following long-term ovariectomy with no estradiol exposure, there was an effect of treatment (p<0.05) on hippocampal ERα (A) and aromatase (B) expression, with post hoc testing revealing increased levels of both proteins in the JB1 group as compared to the Veh group. Following long-term ovariectomy with previous estradiol exposure, there was an effect of treatment (p<0.05) on hippocampal ERα (C) and aromatase (D) expression with post hoc testing revealing decreased levels in the JB1, Let, and JB1+Let groups as compared to the Veh group. *p<0.05 vs. Veh

There was a main effect of treatment on hippocampal aromatase levels, as shown in Figure 9B (F(3,37)= 6.65, p=0.001). Post hoc testing revealed a significant increase in aromatase expression in the JB1 group (p=0.012) as compared to the Veh group. There was no difference in aromatase levels between the Veh group and the Let group (p=0.974) or between the Veh group and the JB1+Let group (p=0.403).

*Neither antagonism of IGF-1 receptors nor inhibition of neuroestrogen synthesis impacted levels of locally synthesized neuroestrogens. The lack of effect of letrozole on estradiol levels suggests low baseline levels of locally synthesized neuroestrogens in the hippocampus following long-term ovariectomy*.

After chronic treatment with either JB1, letrozole, or JB1 plus letrozole following long-term ovariectomy, there was no effect of treatment on hippocampal estradiol levels (Figure 10A; F(3,29)= 0.466, p=0.708).

**Figure 10.**
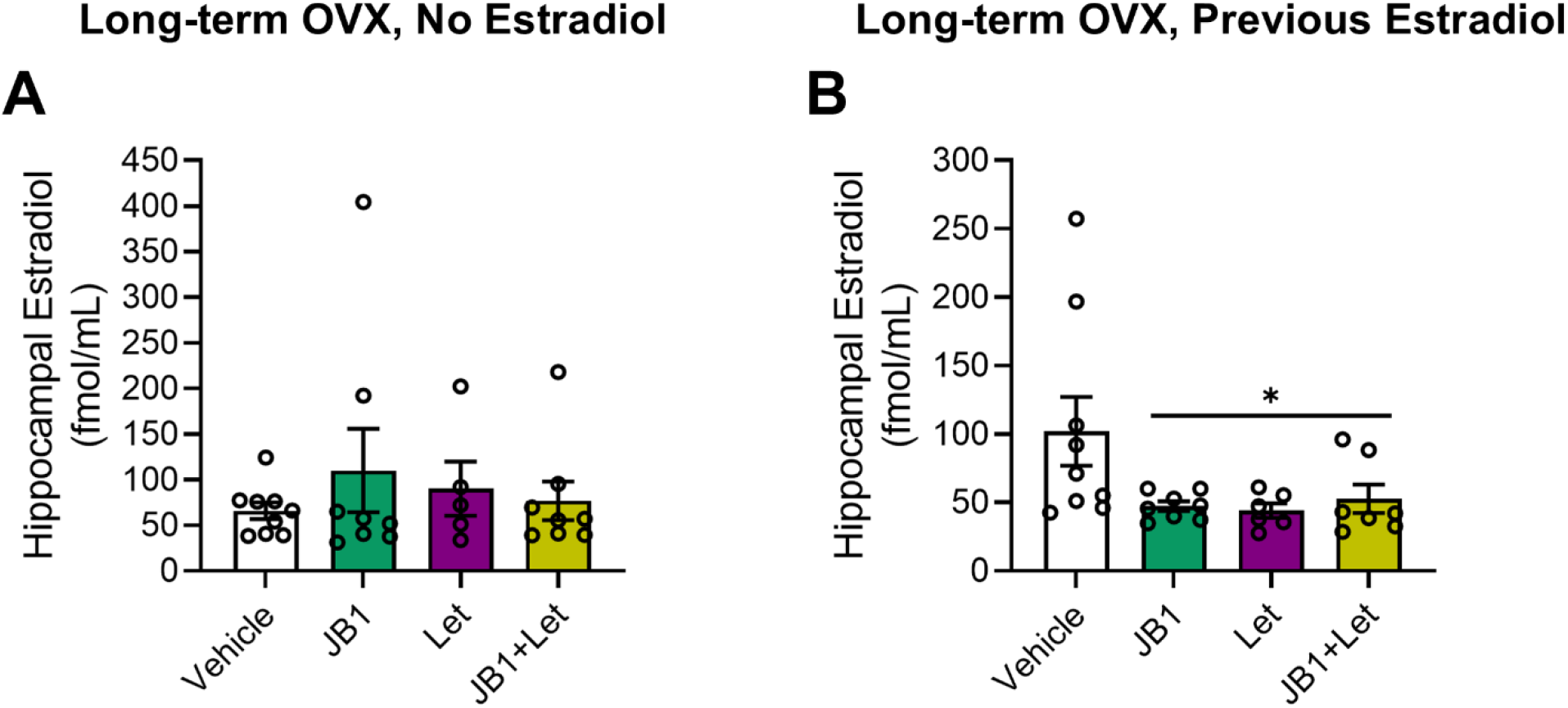
Impacts of chronic JB1, Letrozole, or JB1+Letrozole treatment on hippocampal estradiol levels in long-term ovariectomized rats with or without previous midlife estradiol exposure. Middle-aged female rats were ovariectomized and immediately implanted with vehicle (Long-term OVX, No Estradiol groups) or estradiol (Long-term OVX, Previous Estradiol groups) capsules. Forty days later, capsules were removed. One-hundred days following loss of circulating estrogens (either via ovariectomy in Long-term OVX, No Estradiol group or removal of estradiol capsule in Long-term OVX, Previous Estradiol group), rats were trained on the 8-arm radial maze task and subsequently treated with chronic i.c.v. administration of either vehicle (Veh), the IGF-1R antagonist JB1 (JB1), the aromatase inhibitor letrozole (Let), or JB1 and letrozole (JB1+Let). After rats were tested on the radial-arm maze, hippocampi were dissected and processed for estradiol detection via UPLC-MS/MS. Estradiol levels are expressed in fmol/mL. A) Following long-term ovariectomy with no estradiol exposure, there was no effect of treatment on hippocampal estradiol levels. B) Following long-term ovariectomy with previous estradiol exposure, there as an effect of treatment (p<0.05) on estradiol levels, with post hoc testing revealed decreased levels in the JB1, Let, and JB1+Let groups as compared to the Veh group. *p<0.05 vs. Veh

### Experiment 4: A history of previous midlife estradiol treatment protects hippocampal function and memory following long-term ovarian hormone deprivation by maintaining the interactive relationship between IGF-1 and neuroestrogen signaling

Results of Experiment 3 revealed that long-term ovarian deprivation disrupts the ability of IGF-1 and neuroestrogens to exert positive interactive effects on memory as indicated by an enhancement resulting from IGF-1 receptor antagonism and a lack of disruptive effects on memory of letrozole treatment alone. In contrast to the Experiment 3 results in which JB1 enhanced memory, previous work from our lab revealed that antagonism of IGF-1 receptors by JB1 disrupts memory in ovariectomized rats treated with ongoing (Nelson et al. 2014) or previous (Witty et al. 2013) estradiol. In Experiment 4, we tested the hypothesis that previous midlife estradiol exposure maintains the positive impact of IGF-1 signaling on the MAPK signaling pathway, ERα levels and subsequent impact on memory by sustaining neuroestrogen synthesis in the hippocampus long-term, even after termination of estradiol treatment and in the absence of circulating estrogens. We employed a rat model of menopause in which middle-aged animals received an estradiol implant at the time of ovariectomy that was removed following 40 days of treatment, modeling women who take hormones for a few years and then stop. One hundred days following termination of estradiol treatment, rats were treated with chronic icv delivery of vehicle (Veh group), IGF-1R antagonist JB1 (JB1 group), aromatase inhibitor letrozole (Let group), or JB1 plus letrozole (JB1+Let group). Rats were then tested on a hippocampal-dependent spatial memory task and later hippocampal levels of MAPK and PI3K-Akt pathway activation, ERα and aromatase expression, and estradiol levels were measured.

*Antagonizing IGF-1 receptor activity, inhibition of neuroestrogen synthesis, or the combination of both exert similar detrimental effects on spatial memory. Results indicate that a previous history of estradiol treatment allows for long-term maintenance of the beneficial interactive effects of IGF-1 and neuroestrogens in the hippocampus, in which both are necessary, but neither sufficient to enhance memory*.

As illustrated in 7, Mixed-Design ANOVA revealed a main effect of delay (F(4,124)=20.720; p=4.26×10^-13^) and a main effect of treatment (F(3,31)=3.205; p=0.037) on radial-arm maze performance. Post hoc testing revealed significantly more errors of 8 across delays in the JB1 group (p=0.033), the Let group (p=0.033), and the JB1+Let group (p=0.012) as compared to the Veh group. There was a statistically trending interaction between delay and treatment (F(12, 124)=1.655; p=0.085).

*Antagonizing IGF-1 receptor activity, inhibition of neuroestrogen synthesis, or the combination of both, resulted in similar decreased levels of MAPK activation, and no effects of on Akt activation. Results indicate that after a previous history of estradiol treatment, MAPK signaling pathway predominates due to interactions of IGF-1 and neuroestrogen signaling*.

After chronic treatment with either JB1, letrozole, or JB1 plus letrozole following previous estradiol exposure, there was an effect of treatment on phosphorylation of both p44-MAPK (Figure 8D; F(3,34)= 3.694, p=0.022) and p42-MAPK (Figure 8E; F(3,34)= 4.839, p=0.007). Post hoc testing revealed a significant decrease in phosphorylation of both phosphorylation sites of MAPK in the JB1 group (p44-MAPK, p=0.022; p42-MAPK, p=0.008), the Let group (p44-MAPK, p=0.035; p42-MAPK, p=0.007), and the JB1+Let group (p44-MAPK, p=0.034; p42-MAPK, p=0.038) as compared to the Veh group.

As illustrated in Figure 8F, there was an effect of treatment on phosphorylation of Akt (F(3,34)= 3.248, p=0.035). However, post hoc testing revealed no significant differences between the Veh group and the JB1 (p=0.363), Let (p=0.350), or JB1+Let (p=0.615) groups.

*Antagonizing IGF-1 receptor activity, inhibition of neuroestrogen synthesis, or the combination of both results in similar decreases in protein levels of ERα and aromatase in the hippocampus*.

As illustrated in Figure 9C, there was an effect of treatment on hippocampal ERα levels (F(3,34)= 4.008, p=0.016). Post hoc testing revealed a significant decrease in ERα expression in the JB1 group (p=0.034), the Let group (p=0.023), and the JB1+Let group(p=0.016) as compared to the Veh group.

Finally, there was an effect of treatment on hippocampal aromatase levels, as shown in Figure 9D (F(3,34)= 8.803, p=2.27×10^-4^). Post hoc testing revealed a significant decrease in aromatase expression in the JB1 group (p=0.009), the Let group (p=4.43×10^-4^), and the JB1+Let group (p=2.26×10^-4^) as compared to the Veh group.

*Antagonizing IGF-1 receptor activity, inhibition of neuroestrogen synthesis, or the combination of both results in similar decreased levels of locally synthesized neuroestrogens. The effect of letrozole on estradiol levels suggests that local synthesis of neuroestrogens in the hippocampus is maintained by a history of previous midlife estradiol treatment following long-term ovariectomy*.

After chronic treatment with either JB1, letrozole, or JB1 plus letrozole following previous estradiol exposure, there was an effect of treatment on hippocampal estradiol levels (Figure 10B; F(3,29)=3.10, p=0.044). Post hoc testing revealed a significant decrease in estradiol expression in the JB1 group (p=0.024), the Let group (p=0.027), and the JB1+Let group (p=0.048) as compared to the Veh group.

## Discussion

Results reveal that short-term estrogen treatment during midlife—as is commonly used during the menopausal transition in humans—provides lasting benefits for hippocampal function and memory by robustly altering the interactive relationship between insulin-like growth factor-1 (IGF-1) and locally synthesized neuroestrogens in mediating ligand-independent activation of hippocampal estrogen receptor (ER) α. First, we showed in recently ovariectomized rats (∼10-day) that neuroestrogen synthesis is required for IGF-1-mediated increases in phosphorylation of ERα, activation of the MAPK pathway, and enhanced performance on the hippocampal-dependent radial-arm maze. Next, we found that following long-term ovariectomy (∼100-day), IGF-1 signaling and neuroestrogen signaling no longer provided the same benefits for hippocampal function and memory, demonstrating a weakened relationship between the two hormones following long periods of ovarian hormone deprivation. Remarkably, short-term (40-day) treatment with estradiol immediately following ovariectomy successfully maintained the relationship between IGF-1 and neuroestrogen signaling, resulting in enhanced memory, increased hippocampal activation of MAPK, protein expression of ERα and aromatase, and estradiol levels. Together, results provide a potential model for combatting postmenopausal cognitive decline in which short-term estradiol treatment near the loss of ovarian hormones can sustain hippocampal function and memory by maintaining the dynamic relationships between ERα, IGF-1R, and neuroestrogen synthesis in the aging female brain.

### Effects of IGF-1 on ERα activation, MAPK signaling and memory rely on local estrogen production

Results of Experiment 1 revealed that infusion of IGF-1 to brains of ovariectomized rats increased phosphorylation of hippocampal ERα at S118—a site associated with decreased degradation (Valley et al. 2005) and increased transcriptional activity (Duterte and Smith, 2003) of the receptor. Subcellular compartment fractionation allowed us to localize the increased pS118-ERα that occurred one hour following IGF-1 infusion to the cytosolic compartment of hippocampal cells. Results are consistent with *in vitro* work demonstrating peak dimerization (and presumably therefore, nuclear translocation) of ERα does not occur until two hours after estrogen treatment (Powell and Xu, 2008). Twenty-four hours after infusion of IGF-1, overall ERα levels were increased in the nuclear compartment of hippocampal cells. Results suggest that IGF-1 activation of ERα via phosphorylation at S118 promotes nuclear translocation of ERα, protecting the receptor from degradation and allowing for sustained ERα levels. Furthermore, results implicate a role for locally synthesized neuroestrogens in IGF-1 effects. Inhibition of local synthesis of neuroestrogens via administration of letrozole blocked the ability of IGF-1 to increase phosphorylation of ERα and the subsequent increase in nuclear ERα protein levels.

A potential mechanism by which IGF-1 and neuroestrogen interact to impact ERα is via intracellular signaling pathways. Both MAPK and PI3K-Akt signaling are activated via tyrosine kinase receptor IGF-1 or by neuroestrogens acting on membrane-bound estrogen receptors (Russo et al. 2005; Foster 2012). Here, infusions of IGF-1, but not IGF-1 plus letrozole, increase phosphorylation of p44- and p42-MAPK. Letrozole had no impact on IGF-1-induced increase in Akt phosphorylation. Earlier work in cell culture demonstrated that MAPK phosphorylates ERα at S118 following IGF-1 treatment (Kato et al. 1995), and recent *in vitro* work from our lab support the role of neuroestrogens in activating the MAPK pathway in conjunction with IGF-1R (Pollard and Daniel, 2019). Interestingly, Pollard and Daniel (2019) also demonstrated a mutually repressive relationship between MAPK and Akt in which both pathways inhibit each other, allowing for highly regulated control of ERα activity by IGF-1R. In summary, data indicate that IGF-1 and neuroestrogen signaling interact via the MAPK, but not the Akt, pathway, to activate hippocampal ERα in the absence of circulating estrogens. The significance of these interactions is supported by the results of Experiment 2 in which IGF-1 mediated enhancement of a hippocampal dependent radial-maze task was blocked by letrozole, indicating that IGF-1 activation of ERα requires neuroestrogen synthesis to enhance hippocampal memory.

### Effects of IGF-1 on MAPK signaling and memory are significantly altered as a result of long-term loss of ovarian function and associated putative decrease in neuroestrogens

Local inhibition of aromatase activity has been shown to impair memory consolidation in recently ovariectomized mice (Tuscher et al 2016). However, hippocampal aromatase expression (Ma et al 2020), estradiol levels (Chen et al 2021), and neuroestrogen-mediated transcriptional activity (Baumgartner et al 2019) decrease following long-term, but not short-term, ovariectomy. Consistent with those findings are results of Experiment 3 in which blocking neuroestrogen synthesis via letrozole administration had no impact on hippocampal memory in long-term ovariectomized animals. Surprisingly, pharmacologically inhibiting IGF-1R using JB1 actually enhanced memory in long-term ovariectomized rats, suggesting the possibility that IGF-1 signaling becomes detrimental following long-term ovarian hormone deprivation and associated loss of local synthesis of neuroestrogens.

The paradoxical beneficial effect of IGF-1 antagonism on memory following long-term ovarian hormone deprivation could potentially be explained by regulation of aromatase activity via IGF-1 signaling. In addition to enhancement of memory, blocking IGF-1R with JB1 in long-term ovariectomized animals resulted in increased hippocampal MAPK activation, decreased PI3K-Akt activation, and increased expression of ERα and aromatase. Importantly, JB1 plus letrozole did not have the same effects on memory and protein expression as JB1 administered alone, indicating that the positive impacts of JB1 on memory require subsequent neuroestrogens synthesis. While the precise mechanism for activation of the enzyme aromatase is far from clear—with certain phospho-sites associated with increased activity, and others associated with suppressed activity (Balthazart et al. 2005; Catalano et al. 2009; Miller et al. 2008)—its activation can be regulated by kinase cascades initiated by IGF-1R and membrane estrogen receptors. For example, in T47D breast cancer cells, inhibition of the Akt pathway was associated with increased aromatase activity (Su et al 2011). Here, results suggest a mechanism in which inhibiting IGF-1R results in decreased PI3K-Akt activation, which in turn disinhibits the MAPK pathway and allows for increased ERα and aromatase. Ultimately, however, we detected no group differences in hippocampal estradiol levels following long-term ovariectomy, likely due to overall decreases in estradiol levels following long periods of ovarian hormone deprivation reported previously (Chen et al 2021). Nevertheless, results suggest that a shift in IGF-1 signaling from MAPK to PI3K-Akt following long-term ovarian hormone deprivation is detrimental to memory.

### Effects of long-term loss of ovarian function are mitigated by early, short-term estrogen treatment, and reflect effects on levels of aromatase and neuroestrogens

Results of Experiment 4 demonstrate that a history of previous estradiol treatment reverses the negative effects of IGF-1 signaling on the hippocampus and memory in long-term ovariectomized rats. Consistent with earlier work (Witty et al. 2013), we found that JB1 treatment impaired memory and decreased levels of MAPK phosphorylation and ERα expression in animals previously treated with estradiol during midlife. Here we extend those findings by demonstrating the necessary role for neuroestrogens in facilitating activation of the MAPK pathway by IGF-1R. We found identical effects of JB1, letrozole, and JB1 plus letrozole on memory, MAPK phosphorylation, ERα and aromatase protein expression, and hippocampal estradiol levels in animals that experienced previous estradiol treatment, indicating that IGF-1R and neuroestrogens work together to maintain hippocampal function in aging females following a previous period of midlife estradiol treatment.

The current results reveal diverging paths for hippocampal function in two models of menopause. On one path, long-term loss of ovarian hormones results in decreased neuroestrogen activity, shifting the balance of IGF-1 signaling such that activation of the Akt pathway predominates over activation of the MAPK pathway, and leading to decreases in levels of aromatase and phosphorylation of ERα. On the other path, a short-term period of estradiol treatment immediately following loss of ovarian function reverses the negative impact of long-term hormone deprivation on hippocampal function by sustaining levels of neuroestrogens well beyond the period of estradiol treatment, allowing for IGF-1 mediated activation of the MAPK pathway to predominate over the Akt pathway. MAPK signaling leads to increased aromatase expression, continued neuroestrogens synthesis, and phosphorylation of ERα at phospho-site Ser-118. This activation via ligand-independent mechanisms results in dimerization and nuclear translocation of ERα, allowing for sustained levels of the receptor and leading to transcriptional changes that impact hippocampal function and ultimately enhance memory.

## CONCLUSIONS

Collectively, results indicate that short-term estrogen treatment following midlife loss of ovarian function has long-lasting effects on hippocampal function and memory by dynamically regulating cellular mechanisms that promote activity of ERα in the absence of circulating estrogens. Findings demonstrate how changes in hippocampal ERα expression, IGF-1R signaling, and neuroestrogen synthesis following long-term ovariectomy can negatively impact memory, but that a history of previous estradiol treatment protects the hippocampus against these changes to combat cognitive decline in rodent models of menopause.

## Acknowledgements

This work was supported by the National Institute on Aging Grant RF1AG041374 to JMD. In addition, this project used the services of the University of Pittsburgh Small Molecule Biomarker Core, which was partially funded by NIH through S10RR023461 and S10OD028540.

## References

Balthazart J, Baillien M, Ball G. Interactions Between Kinases and Phosphatases in the Rapid Control of Brain Aromatase. J Neuroendocrinol. 2005 Sep;17(9), 553–559. doi: 10.1111/j.1365-2826.2005.01344.x

Baumgartner NE, Black KL, McQuillen SM, Daniel JM. (2021). Previous estradiol treatment during midlife maintains transcriptional regulation of memory-related proteins by ERα in the hippocampus in a rat model of menopause. Neurobiol Aging. 2021 Sep;105:365–373. doi: 10.1016/j.neurobiolaging.2021.05.022. Epub 2021 Jun 5. PMID: 34198140; PMCID: PMC8338908.

Baumgartner NE, Daniel JM. Estrogen receptor α: a critical role in successful female cognitive aging. Climacteric. 2021 Aug;24(4):333–339. doi: 10.1080/13697137.2021.1875426. Epub 2021 Feb 1. PMID: 33522313; PMCID: PMC8273070.

Baumgartner NE, Grissom EM, Pollard KJ, McQuillen SM, Daniel JM. Neuroestrogen-Dependent Transcriptional Activity in the Brains of ERE-Luciferase Reporter Mice following Short- and Long-Term Ovariectomy. eNeuro. 2019 Oct 16;6(5):ENEURO.0275-19.2019. doi: 10.1523/ENEURO.0275-19.2019. PMID: 31575604; PMCID: PMC6795557.

Black KL, Witty CF, Daniel JM. Previous Midlife Oestradiol Treatment Results in Long-Term Maintenance of Hippocampal Oestrogen Receptor α Levels in Ovariectomised Rats: Mechanisms and Implications for Memory. J Neuroendocrinol. 2016 Oct;28(10):10.1111/jne.12429. doi: 10.1111/jne.12429. PMID: 27603028; PMCID: PMC5527336.

Cardona-Gómez GP, DonCarlos L, Garcia-Segura LM. Insulin-like growth factor I receptors and estrogen receptors colocalize in female rat brain. Neuroscience. 2000;99(4):751–60. doi: 10.1016/s0306-4522(00)00228-1. PMID: 10974438.

Catalano S, Barone I, Giordano C, Rizza P, Qi H, Gu G, Malivindi R, Bonofiglio D, Andò S. Rapid estradiol/ERalpha signaling enhances aromatase enzymatic activity in breast cancer cells. Mol Endocrinol. 2009 Oct;23(10), 1634–1645. doi: 10.1210/me.2009-0039

Chen HB, Xu C, Zhou MH, Qiao H, An SC. Endogenous hippocampal, not peripheral, estradiol is the key factor affecting the novel object recognition abilities of female rats. Behav Neurosci. 2021 Oct;135(5):668–679. doi: 10.1037/bne0000480. Epub 2021 Aug 16. PMID: 34398621.

Daniel J.M. (2015) The Land Radial-Arm Maze: Eight Out of Eight Arms Baited with Food Protocol for Rodents. In: Bimonte-Nelson H. (eds) The Maze Book. Neuromethods, vol 94. Humana Press, New York, NY. https://doi.org/10.1007/978-1-4939-2159-1_17

Dutertre M, Smith CL. Ligand-independent interactions of p160/steroid receptor coactivators and CREB-binding protein (CBP) with estrogen receptor-alpha: regulation by phosphorylation sites in the A/B region depends on other receptor domains. Mol Endocrinol. 2003 Jul;17(7):1296–314. doi: 10.1210/me.2001-0316. Epub 2003 Apr 24. PMID: 12714702.

Foster TC. Role of estrogen receptor alpha and beta expression and signaling on cognitive function during aging. Hippocampus. 2012 Apr;22(4):656–69. doi: 10.1002/hipo.20935. Epub 2011 Apr 27. PMID: 21538657; PMCID: PMC3704216.

Grissom EM, Daniel JM. Evidence for Ligand-Independent Activation of Hippocampal Estrogen Receptor-α by IGF-1 in Hippocampus of Ovariectomized Rats. Endocrinology. 2016 Aug;157(8):3149–56. doi: 10.1210/en.2016-1197. Epub 2016 Jun 2. PMID: 27254005; PMCID: PMC4967122.

Henderson VW, Watt L, Buckwalter JG. Cognitive skills associated with estrogen replacement in women with Alzheimer’s disease. Psychoneuroendocrinology. 1996 May;21(4):421–30. doi: 10.1016/0306-4530(95)00060-7. PMID: 8844880.

Kato S, Endoh H, Masuhiro Y, Kitamoto T, Uchiyama S, Sasaki H, Masushige S, Gotoh Y, Nishida E, Kawashima H, Metzger D, Chambon P. Activation of the estrogen receptor through phosphorylation by mitogen-activated protein kinase. Science. 1995 Dec 1;270(5241):1491–4. doi: 10.1126/science.270.5241.1491. PMID: 7491495.

Li J, Gibbs RB. Detection of estradiol in rat brain tissues: Contribution of local versus systemic production. Psychoneuroendocrinology. 2019 Apr;102:84–94. doi: 10.1016/j.psyneuen.2018.11.037. Epub 2018 Dec 1. PMID: 30529907.

Li J, Oberly PJ, Poloyac SM, Gibbs RB. A microsomal based method to detect aromatase activity in different brain regions of the rat using ultra performance liquid chromatography-mass spectrometry. J Steroid Biochem Mol Biol. 2016 Oct;163:113–20. doi: 10.1016/j.jsbmb.2016.04.013. Epub 2016 Apr 22. PMID: 27113434.

Ma Y, Liu M, Yang L, Zhang L, Guo H, Hou W, Qin P. Loss of Estrogen Efficacy Against Hippocampus Damage in Long-Term OVX Mice Is Related to the Reduction of Hippocampus Local Estrogen Production and Estrogen Receptor Degradation. Mol Neurobiol. 2020 Aug;57(8):3540–3551. doi: 10.1007/s12035-020-01960-z. Epub 2020 Jun 15. PMID: 32542593.

Mendez P, Azcoitia I, Garcia-Segura LM. Estrogen receptor alpha forms estrogen-dependent multimolecular complexes with insulin-like growth factor receptor and phosphatidylinositol 3-kinase in the adult rat brain. Brain Res Mol Brain Res. 2003 Apr 10;112(1-2):170–6. doi: 10.1016/s0169-328x(03)00088-3. PMID: 12670715.

Miller TW, Shin I, Kagawa N, Evans DB, Waterman MR, Arteaga CL. Aromatase is phosphorylated in situ at Serine-118. J Steroid Biochem Mol Biol. 2008 Nov;112(1-3), 95–101. doi: 10.1016/j.jsbmb.2008.09.001

Nelson BS, Black KL, Daniel JM. Circulating Estradiol Regulates Brain-Derived Estradiol via Actions at GnRH Receptors to Impact Memory in Ovariectomized Rats. eNeuro. 2016 Dec 13;3(6):ENEURO.0321-16.2016. doi: 10.1523/ENEURO.0321-16.2016. PMID: 28032117; PMCID: PMC5172373.

Nelson BS, Springer RC, Daniel JM. Antagonism of brain insulin-like growth factor-1 receptors blocks estradiol effects on memory and levels of hippocampal synaptic proteins in ovariectomized rats. Psychopharmacology. 2014 Mar; 231(5):899–907. doi: 10.1007/s00213-013-3310-7. PMID: 24146138.

Pollard KJ, Daniel JM. Nuclear estrogen receptor activation by insulin-like growth factor-1 in Neuro-2A neuroblastoma cells requires endogenous estrogen synthesis and is mediated by mutually repressive MAPK and PI3K cascades. Mol Cell Endocrinol. 2019 Jun 15;490:68–79. doi: 10.1016/j.mce.2019.04.007. Epub 2019 Apr 13. PMID: 30986444; PMCID: PMC6520186.

Pollard KJ, Wartman HD, Daniel JM. Previous estradiol treatment in ovariectomized mice provides lasting enhancement of memory and brain estrogen receptor activity. Horm Behav. 2018 Jun;102:76–84. doi: 10.1016/j.yhbeh.2018.05.002. Epub 2018 May 12. PMID: 29742445; PMCID: PMC6004337.

Powell E, Wang Y, Shapiro DJ, Xu W. Differential requirements of Hsp90 and DNA for the formation of estrogen receptor homodimers and heterodimers. J Biol Chem. 2010 May 21;285(21):16125–34. doi: 10.1074/jbc.M110.104356. Epub 2010 Mar 30. PMID: 20353944; PMCID: PMC2871481.

Rodgers SP, Bohacek J, Daniel JM. Transient estradiol exposure during middle age in ovariectomized rats exerts lasting effects on cognitive function and the hippocampus. Endocrinology. 2010 Mar;151(3):1194–203. doi: 10.1210/en.2009-1245. Epub 2010 Jan 12. PMID: 20068005.

Russo VC, Gluckman PD, Feldman EL, Werther GA. The insulin-like growth factor system and its pleiotropic functions in brain. Endocr Rev. 2005 Dec;26(7):916–43. doi: 10.1210/er.2004-0024. Epub 2005 Aug 30. PMID: 16131630.

Sherwin BB. Estrogenic effects on memory in women. Ann N Y Acad Sci. 1994 Nov 14;743:213–30; discussion 230-1. doi: 10.1111/j.1749-6632.1994.tb55794.x. PMID: 7802415.

Sohrabji F. Estrogen-IGF-1 interactions in neuroprotection: ischemic stroke as a case study. Front Neuroendocrinol. 2015 Jan;36:1–14. doi: 10.1016/j.yfrne.2014.05.003.

Su B, Wong C, Hong Y, Chen S. Growth factor signaling enhances aromatase activity of breast cancer cells via post-transcriptional mechanisms. J Steroid Biochem Mol Biol. 2011 Feb;123(3-5), 101–108. doi: 10.1016/j.jsbmb.2010.11.012

Tuscher JJ, Szinte JS, Starrett JR, Krentzel AA, Fortress AM, Remage-Healey L, Frick KM. Inhibition of local estrogen synthesis in the hippocampus impairs hippocampal memory consolidation in ovariectomized female mice. Horm Behav. 2016 Jul;83, 60–67. doi: 10.1016/j.yhbeh.2016.05.001

Valley CC, Métivier R, Solodin NM, Fowler AM, Mashek MT, Hill L, Alarid ET. Differential regulation of estrogen-inducible proteolysis and transcription by the estrogen receptor alpha N terminus. Mol Cell Biol. 2005 Jul;25(13):5417–28. doi: 10.1128/MCB.25.13.5417-5428.2005. PMID: 15964799; PMCID: PMC1156995.

Witty CF, Gardella LP, Perez MC, Daniel JM. Short-term estradiol administration in aging ovariectomized rats provides lasting benefits for memory and the hippocampus: a role for insulin-like growth factor-I. Endocrinology. 2013 Feb;154(2):842–52. doi: 10.1210/en.2012-1698. Epub 2012 Dec 21. PMID: 23264616.

